# Exploiting Epigenetic Targets to Overcome Taxane Resistance in Prostate Cancer

**DOI:** 10.1101/2023.08.10.552560

**Authors:** Buse Cevatemre, Ipek Bulut, Beyza Dedeoglu, Arda Isiklar, Hamzah Syed, Ozlem Yedier Bayram, Tugba Bagci-Onder, Ceyda Acilan Ayhan

## Abstract

The development of taxane resistance remains a major challenge for castration resistant prostate cancer (CR-PCa), despite the effectiveness of taxanes in prolonging patient survival. To uncover novel targets, we performed an epigenetic drug screen on taxane (docetaxel and cabazitaxel) resistant CR-PCa cells. We identified BRPF reader proteins, along with several epigenetic groups (CBP/p300, Menin-MLL, PRMT5 and SIRT1) that act as targets effectively reversing the resistance mediated by ABCB1. Targeting BRPFs specifically resulted in the resensitization of resistant cells, while no such effect was observed on the sensitive compartment. These cells were successfully arrested at the G_2_/M phase of cell cycle and underwent apoptosis upon BRPF inhibition, confirming the restoration of taxane susceptibility. Pharmacological inhibition of BRPFs reduced ABCB1 activity, indicating that BRPFs may be involved in an efflux-related mechanism. Indeed, ChIP-qPCR analysis confirmed binding of BRPF1 to the ABCB1 promoter suggesting direct regulation of the ABCB1 gene at the transcriptional level. RNA-seq analysis revealed that BRPF1 knockdown affects the genes enriched in mTORC1 and UPR signaling pathways, revealing potential mechanisms underlying its functional impact, which is further supported by the enhancement of taxane response through the combined inhibition of ABCB1 and mTOR pathways, providing evidence for the involvement of multiple BRPF1-regulated pathways. Beyond clinical attributes (Gleason score, tumor stage, therapy outcome, recurrence), metastatic PCa databases further supported the significance of BRPF1 in taxane resistance, as evidenced by its upregulation in taxane-exposed PCa patients.

## Introduction

Prostate cancer (PCa) is the second most common malignancy among men worldwide, comprising 13.5% of all male cancer diagnoses (Bray et al., 2018). In early stages of the disease, prostatectomy and radiotherapy are considered as the main therapeutic options. However, 20–30% of patients relapse after 5-10 years, and Androgen Deprivation Therapy (ADT) is currently the primary approach for advanced disease. Although the majority of patients initially respond to ADT, PCa relapse is frequently observed in a few years leading to progression to the castration resistant PCa (CR-PCa) stage (Hamberg et al., 2008; Sternberg, 2008). The major treatment option for metastatic CR-PCa includes the use of taxanes (mainly docetaxel, followed by cabazitaxel). Nevertheless, despite the prolonged survival resulting from docetaxel, most patients become refractory due to the development of resistance and succumb to PCa. Thus, it is crucial to find alternative options to overcome resistance and resensitize patients to taxanes.

One of the major factors that contribute to drug resistance is alterations in the epigenome of cancer cells. Not surprisingly, epigenetic modifiers have been targeted by numerous researchers, and several clinical trials are currently recruiting patients to study the efficacy of epigenetic modifiers in PCa, including but not limited to inhibitors of bromodomain and extra-terminal domain (BET), histone methyltransferases (HMT), DNA methyltransferases (DNMT), histone deacetylases (HDAC) or CBP/p300 (Ge et al., 2020; Ponnusamy et al., 2020; Kumaraswamy et al., 2021). More importantly, some of these studies test epi-drugs in CR-PCa and progressive patients either as monotherapy or in combination with other drugs, including Dtx (López et al., 2022; Liao and Xu, 2019). These studies show that epigenetic drugs have the potential to overcome chemotherapy resistance in tumors.

Taxane resistance has been extensively investigated, revealing multiple mechanisms involved, such as upregulated taxane-metabolizing enzymes, prosurvival pathways, altered microtubule regulatory proteins, EMT induction, and dysregulated non-coding RNAs, among other identified contributors (Sekino et al., 2020; Lombard et al., 2020; Maloney et al., 2020; Mosca et al., 2021). Despite the involvement of multiple pathways in the resistance phenotype, the major underlying factor is the drug efflux mediated by the ABCB1 transporter.

In this study, we established two Dtx- and Cbz-resistant CR-PCa cells (Du145 and 22Rv1) and initially characterized the transcriptomic changes in these cells using RNA-seq analysis. Indeed, ABCB1 was one of the top hits, whose expression was greatly elevated. Interestingly, there was no increase in the copy number of ABCB1 gene in 3 out of 4 resistant cell lines, indicating that the transcriptional activity of ABCB1 was altered during the resistance process. Therefore, under the light of new developments in the PCa continuum, we assessed the role of epigenetic modifiers for their efficacy to revert taxane resistance in CR-PCa and conducted an epi-drug screen to uncover the resensitizing factors. We were able to identify five different groups of modulators (CBP/p300, Menin-MLL, SIRT, PRMT5 and BRPF) that were able to effectively revert resistance, all of which were verified via further analyses. Of these, BRPF inhibitors were essential for the resistant, but not for the sensitive parental cells. Silencing BRPF proteins showed a similar phenotype, supporting the importance of BRPFs for resistance. BRPF inhibition reduced ABCB1 activity and gene expression, suggesting the involvement of BRPFs in drug efflux. We observed the occupancy of the ABCB1 promoter by BRPF1 through ChIP-qPCR analysis, providing strong evidence for the direct involvement of BRPF1 in the regulation of the ABCB1 gene. By performing RNA-seq analysis following BRPF inhibition, we identified the modulation of genes associated with the mTORC1 and UPR signaling pathways. Silencing these pathways mimicked BRPF inhibition and reverted resistance, potentially explaining how BRPFs may overcome taxane resistance. Furthermore, the co-targeting of ABCB1 and mTOR pathways enhanced the response to taxanes, providing evidence for the involvement of multiple pathways under the regulation of BRPF1. Lastly, our analysis of BRPF1 expression in pan-cancer and PCa databases demonstrated its significant association with various clinical attributes, including the upregulation observed in taxane-exposed PCa patients, suggesting BRPF1 as a potential biomarker for the progression of PCa.

## Materials and Methods

### Generation of Taxane-Resistant Prostate Cancer Cell Models

Du145 (ATCC no. HTB-81) and 22Rv1 (ATCC no. CRL-2505) cells were maintained in RPMI-1640 medium (Gibco, 11875093) supplemented with 10% fetal bovine serum (Biowest, S1810) and 1% penicillin/streptomycin (Biowest, L0022). Cells were grown at 37°C in a humidified atmosphere with 5% CO_2_. Resistant clones were selected by culturing cells with docetaxel (Sigma-Aldrich, 01885) and/or cabazitaxel (Sigma-Aldrich, SML2487) for 72 h. A dose-escalation strategy was implemented, beginning at 1 nM, and doubled to reach 125 nM. Parental cells were passaged alongside as an age-matched appropriate control.

### CRISPR-Cas9

gRNA sequences were designed using the Benching software (https://www.benchling.com) and are listed in **Sup. Table 1**. LentiCRISPR v2 (Addgene #52961) plasmid DNA (2 µg) was digested with BsmBI-v2 (NEB, R0739) at 55°C for 3 h. The digested vector was purified using the NucleoSpin Gel and PCR Cleanup kit (Macherey-Nagel™, 740609) and then ligated with annealed gRNA oligos. To produce lentivirus, Hek293T cells were co-transfected with packaging and envelope vectors, psPAX2 (Addgene 12260) and pVSVg (Addgene, 14888), respectively. Transduction was carried out using Polybrene (8 µg/ml). Knockout was confirmed by western blotting.

### shRNA Cloning

shRNA sequences were designed using the Broad Institute GPP (https://portals.broadinstitute.org/gpp/public/) and are listed in **Sup. Table 1**. The pLKO.1 (Addgene #8453) lentiviral vector was digested with AgeI (NEB, R3552) and EcoRI (NEB, R3101). Transfection, lentivirus production, and transduction were performed as described for CRISPR-Cas9.

### Cell Proliferation and Doubling Time Analysis

#### Sulforhodamine B (SRB) Viability Assay

Du145 (4 × 10^3^) and 22Rv1 (7.5 × 10^4^) cells were seeded on 96-well plates the day prior to drug exposure. Cells were fixed with 50% (w/v) TCA (Sigma, T6399) at 4°C for 1 h, stained with 0.4% (w/v) SRB (Santa Cruz, sc-253615) for 30 min and subsequently washed with 1% acetic acid. SRB dye was extracted using a 10 mM Trizma base (Sigma, T1503). Measurements were carried out at 564 nm using a microplate reader (Synergy H1 Hybrid reader, BioTek). Epigenetics Screening Library (Cayman, 11076) was used to uncover the epigenetic regulators of taxane resistance. Du145-P/R (4 × 10^3^) and 22Rv1-P/R (1 × 10^4^) cells were co-treated with compounds (5 μM) and IC_25_ values of taxanes for 72 h.

### Colony Formation Assay

Du145 (7.5 × 10^2^) and 22Rv1 (1 × 10^3^) cells were seeded on 12-well plates the day prior to taxane exposure. At the end of the treatment, the medium was replaced by drug-free medium, and the cells were cultured for an additional 10 days. The colonies were fixed for 10 min with methanol (Merck, 1.06009) and stained with 0.5% (w/v) crystal violet (Merck, 1.09218) for 15 min. Quantification was performed using ImageJ (Schneider et al., 2012).

### xCELLigence RTCA

Du145 (5 × 10^3^) and 22Rv1 (1 × 10^4^) cells were seeded on 96-well E-Plates (Roche). Proliferation was monitored every 1 h and time dependent cell index (CI) graphs were generated by using the RTCA software.

### Calcein Retention Assay

Du145 (7.5 × 10^3^) and 22Rv1 (10 × 10^3^) cells were seeded on black 96-well plates the day prior to verapamil (20 µM, 8 h) and BRPF inhibitor exposure (5 µM, 24 h). Cells were then incubated with 125 nM calcein-AM (Invitrogen™, C1430) for 30 min at 37°C. Images were acquired by an inverted fluorescence microscope (Leica DMI8).

### CellTiter-Glo® (CTG) Luminescent Cell Viability Assay

Du145 (4 × 10^3^) and 22Rv1 (7.5 × 10^4^) cells were seeded on 96-well plates the day prior to drug exposure. CellTiter-Glo Reagent (Promega, G7570) was added to the wells (1/10) and incubated for 30 min at 37°C protected from light. Luminescent signal was detected by using a luminometer (Synergy H1 Hybrid reader, BioTek).

### Cell Adhesion Assay

To determine the time required for cells to adhere, Du145-P/R and 22Rv1-P/R cells (1 × 10^6^) were seeded on 24-well plates and allowed to adhere. Unattached cells were washed off with PBS and attached cells were fixed with methanol (Merck, 1.06009) at 20 min intervals up to 6 h after seeding. Attached cells were stained with 0.5% (w/v) crystal violet (Merck, 1.09218). Plates were scanned and particle mean for each well were analyzed using ImageJ.

### Immunofluorescence Staining

Du145-P/R cells grown on coverslips were treated with Dtx or Cbz for 24 h. Cells were fixed with methanol (Merck, 1.06009) for 20 min, permeabilized with 0.1% Triton X-100 at rt for 15 min and blocked with SuperBlock (ScyTek Biotech Life Sciences) at rt for 15 min. Cells were incubated with α-tubulin antibody (Cell Signaling Technology, 76031) at rt for 1 h. Alexa Fluor™ 488 secondary antibody (Thermo Fisher Scientific, A11029) was added and incubated at rt for 1 h. To visualize the nuclei the cells were stained with DAPI (Abcam, ab104139). Images were acquired by a confocal microscope (Leica SP8).

### Cell Cycle Analysis

Du145 (3 × 10^4^) and 22Rv1 (6 × 10^4^) cells were seeded on 12-well plates the day prior to treatments. Cells were washed with PBS and fixed for at least 3 h with 70% ethanol (Merck, 100983). Cells were washed with PBS and analyzed by using the Cell Cycle Kit (Luminex Corporation, MCH100106) according to the manufacturer’s instructions.

### Cell Death Analysis

Du145-DtxR (2 × 10^5^) cells were seeded on 6-well plates the day prior to drug treatments. Cells were harvested by trypsinization and analyzed for cell death mode using Annexin V & Dead Cell Kit (Luminex Corporation, MCH100105) and Caspase-3/7 Kit (Luminex Corporation, MCH100108) according to the manufacturer’s instructions.

### qRt-PCR

Total RNA was extracted with NucleoSpin™ RNA Plus Isolation Kit (Macherey Nagel, 740984.50) and cDNA was synthesized from 1 µg total RNA using the M-MLV Reverse Transcriptase (Thermo Fisher Scientific, 28025013). cDNA was amplified using LightCycler 480 SYBR Green I Master mix (Roche Diagnostics, 04707516001). The reaction mixture contained 0.15 μM specific primers (listed in **Sup. Table 1**) for target genes and incubated at 95°C for 5 min, followed by 40 cycles of 95°C for 10 s, 60°C for 30 s, and 72°C for 1 s. β-Actin was used as reference control and qRt-PCR was run on the PikoReal Real-Time PCR System (Thermo Fisher Scientific). The relative fold change in gene expressions were measured with the 2^(−ΔΔCT)^ method.

### DNA copy numbers by qPCR

Genomic DNA was extracted from 22Rv1 and D145 cells using Nucleospin Tissue kit (Macherey Nagel, 740952.50) according to the manufacturer’s protocol. Using genomic DNA as a template, the quantification of ABCB1 copy number was performed with LightCycler 480 SYBR Green I Master mix (Roche Diagnostics, 04707516001). ABCB1 genomic amplification primer sequences are listed in **Sup. Table 1**.

### SDS-PAGE and Western Blotting (WB)

Proteins were extracted by RIPA lysis buffer (EcoTech Biotechnology, RIPA-100) containing cOmplete™ protease inhibitor cocktail (Merck, 11697498001), PMSF (Merck, 10837091001) and phosSTOP phosphatase inhibitor (Merck, 4906845001) at 4°C for 30 min. Protein concentrations were determined using BCA assay (Thermo Fisher Scientific, 23225). The Laemmli sample buffer (Bio-Rad Laboratories, 1610747) and DL-Dithiothreitol (Sigma, D0632) were added to the proteins and boiled at 95°C for 7 min. Proteins (20-30 µg) were subjected to SDS-PAGE and then transferred to PVDF membranes. The membranes were then blocked with 5% NFDM (Bio-Rad Laboratories, 1706404) at rt for 1 h and blotted with the following primary antibodies at 4°C overnight: β-Actin (Abcam, ab8224), MDR1/ABCB1 (Cell Signaling Technology, 13978), β-Tubulin (Abcam, ab6046), and BRPF1 (Abcam, ab259840). The membranes were washed with TBST and incubated with the HRP-conjugated secondary antibodies (Goat pAb to Rabbit and Mouse IgG, Abcam, ab205718, ab205719) for 1 h at rt. Signal was developed using Immobilon Forte Western HRP substrate (Millipore, WBLUF0100) and imaged by LI-COR Odyssey FC Imaging system (LI-COR Biosciences).

### Cellular Thermal Shift Assay (CETSA)

PCa cells (3.5 × 10^5^) were treated with Dtx (62.5 nM) or Cbz (32.5 nM) for 1 h at 37°C and 5% CO_2_. Treated and untreated cells were centrifuged, and the pellet was resuspended with PBS containing cOmplete™ protease inhibitor cocktail (Merck, 11697498001) and 1 mM PMSF (Merck, 10837091001). The cell suspension then was divided equally and heated for 3 min at indicated temperatures on T100™ Thermal Cycler (Bio-Rad Laboratories). Cells were snap-frozen and thawed twice and centrifuged at 17.000 g for 15 min at 4°C. The protein-containing supernatant was collected and subjected to SDS-PAGE and WB.

### Histone Extraction

Du145 (1 × 10^6^) cells were treated with BRPF inhibitors (5 µM, 1-24h). Treated and untreated cells were lysed and histones were extracted, as previously described (Bulut et al, 2022). The antibodies used were as follows: Acetyl-Histone H3 (Lys9) (Cell Signaling Technologies, 9649); Acetyl-Histone H3 (Lys18) (Cell Signaling Technologies, 13998); Acetyl-Histone H3 (Lys27) (Cell Signaling Technologies, 4353); Histone H3 (Abcam, ab18521).

### RNA-Sequencing and Analysis

Analysis was performed to identify differentially expressed genes; i) between Du145-DtxR and parental cells, ii) Du145-CbzR and parental cells, iii) siBRPF1 and siControl. The comparisons were made with at least 2 biological replicates per treated cell line. Each paired-end sample FASTQ file was assessed for adapter contamination using FastQC. The sequencing data was aligned using STAR (transcriptome mapping) (Dobin et al., 2013), to the GENCODE GRCH38 (Frankish et al., 2019) Human reference transcriptome. Salmon (Patro et al., 2017) was used to quantify the STAR transcriptome BAM files based on the transcripts per million method. In order to aggregate the transcript-level abundance to the gene-level the R package tximport (Soneson et al., 2015) was used. The package DESeq2 (Love et al., 2014) was used to analyze the gene-level count data, identifying differentially expressed genes for each of the three contrasts. Significant differentially expressed genes (DEGs) were defined as genes with a log-fold change > 1 (up-regulated) or log-fold change < -1 (down-regulated) and a false-discovery rate (FDR) adjusted P-value < 0.05. The significant results from the gene expression analysis were annotated using the Biomart database (Durinck et al., 2009), to provide descriptions of gene function. Volcano, MA (base-2 log fold-change versus normalized mean counts), and log-fold change comparison plots generated using ggplot2 were used to visualize the gene expression analysis results.

### ChIP-qPCR

Du145-DtxR cells were crosslinked with %1 formaldehyde (Sigma Aldrich, 818708) at rt for 10 min. Cross-linking was terminated by the addition of glycine (Sigma Aldrich, G7126) (125 mM, 5 min) and cell suspension was centrifuged at 1000 g for 5 min at 4°C. Pellet was washed with cold PBS twice and resuspended in ChIP Lysis Buffer and incubated on ice for 10 min. The released chromatin was sonicated with Bioruptor Plus sonicator (Diagenode) and was centrifuged at 10.000 g for 10 min at 4°C. The supernatant containing the sheared chromatin (fragment sizes of 100-500bp) was collected. The chromatin preparation was incubated with prewashed Dynabeads® (Thermo Fisher protein A magnetic beads, 10001D) at 4°C for 30 min to avoid non-specific binding (preclearing). BRPF1 antibody (Abcam, ab259840) and a non-specific IgG antibody (Sigma Aldrich, PP64B) for negative control were incubated with magnetic beads in PBST for 1 h at rt. The magnetic bead-antibody complex and pre-cleared chromatin preparation was incubated overnight. The magnetic beads were washed with ChIP lysis buffer, low-salt buffer, high salt buffer, LiCl-containing buffer, Tris-EDTA solution twice for 5 min, respectively. ChIP’ed DNA was eluted using an elution buffer containing 1% SDS (Bio-Rad, 1610418) and 0.1M NaHCO_3_ (Merck, S6014) and reverse cross linked by incubating with Rnase A (0.3 mg/ml) (Thermo Fisher Scientific, 12091039) and NaCl (Merck, S9888) (5 M) for 30 min, followed by addition of proteinase K (0.3 mg/ml) (Thermo Fisher Scientific, EO0492) for 1 h. DNA was purified by using the ChIP DNA Clean & Concentrator kit (Zymo Research, D5205). Primers were designed by using UCSC Genome (https://genome.ucsc.edu/) and Ensembl Genome Browser (https://www.ensembl.org/index.html) and primer sequences are listed in **Sup. Table 1**. For the quantitative and simultaneous analysis of immunoprecipitated DNA, qPCR was run on the LightCycler 480 (Roche). ChIP-qPCR results were calculated by % input method.

### Analysis of Clinical Data

Pan-cancer analysis of whole genomes (ICGC/TCGA Pan-Cancer Analysis of Whole Genomes Consortium, 2020), The Cancer Genome Atlas Prostate Adenocarcinoma (TCGA-PRAD) (Abeshouse et al., 2015) and Firehose Legacy, Fred Hutchinson CRC (Kumar et al., 2016), Metastatic Prostate Adenocarcinoma (SU2C/PCF Dream Team, Robinson et al., 2015), Metastatic Prostate Adenocarcinoma (SU2C/PCF Dream Team, Abida et al., 2019), and Metastatic Prostate Adenocarcinoma (MCTP) (Grasso et al., 2012) datasets were accessed via cBioPortal (Cerami et al, 2012; Gao et al., 2013). Statistical analysis using the unpaired t-test was conducted with Prism 8 (GraphPad Software, Inc.).

### Statistical Analysis

All data were analyzed in Prism 8 (GraphPad Software, Inc.) and statistical tests were applied as described in the figure legends. Combination index (CI) values were calculated using CalcuSyn software (Biosoft) based on the method described by Chou and Talalay to assess the synergism (Chou and Talalay, 1984).

## Results

### Establishment and characterization of Dtx and Cbz resistant PCa cells

Dtx and Cbz resistant cell lines (DtxR and CbzR, respectively) were generated from CR-PCa cell lines, Du145 and 22Rv1. The parental cells were allowed to grow for 24 h before being treated (72 h) with dose escalation (**Figure 1A**). Cell growth assays were performed to assess the acquired resistance, revealing a significant number of viable cells in the resistant compartments following taxane exposure **(Figure 1B-C).** The IC_50_ values of resistant cells were significantly higher than those of parental cells exhibiting ∼40-560 fold increase for DtxR and ∼14-170 fold for CbzR **(Table 1).** We showed that these viability differences observed in cells were indeed associated with cell death, as evident by the significant increases in annexin-v and caspase 3/7 positive cells in parental cells (**Figure 1D, Sup. Figure 1A**). Even at higher doses, resistant cells did not display these apoptosis markers, thus reflecting the non-responder state in accordance with the viability assays (**Figure 1D, Sup. Figure 1A).** We proceeded to explore the on-target efficacy of taxanes in both parental and resistant cells. Considering that taxanes exert their activity through microtubule (MT) stabilization, we initially evaluated the MT organization. Taxanes resulted in curved and bundled MTs at the cell periphery in parental lines, while there was no evidence of bundling in the resistant cells (**Figure 1E**). Furthermore, Dtx or Cbz treatments caused a significant G_2_/M arrest in parental cells, whereas the resistant cells were able to bypass the mitotic blockade and re-enter the cell cycle (**Figure 1F, Sup. Figure 1B**). Next, we evaluated the target engagement (TE) by CETSA, a biophysical assay which allows for TE to be measured in intact cells **(Sup. Figure 1C).** As shown in **Figure 1G** and **Sup. Figure 1D**, a higher level of target engagement was observed in parental cells, indicating their ability to retain the drug compared to resistant cells. These findings suggest a potential inadequacy of taxane uptake in resistant cells, leading to a consequent lack of target engagement. In line with our findings, a study involving both sensitive and resistant cells, CETSA TE measurements correlated with taxane sensitivity and efficiently revealed the presence of acquired drug resistance (Langebäck et at., 2019).

**Table 1.**
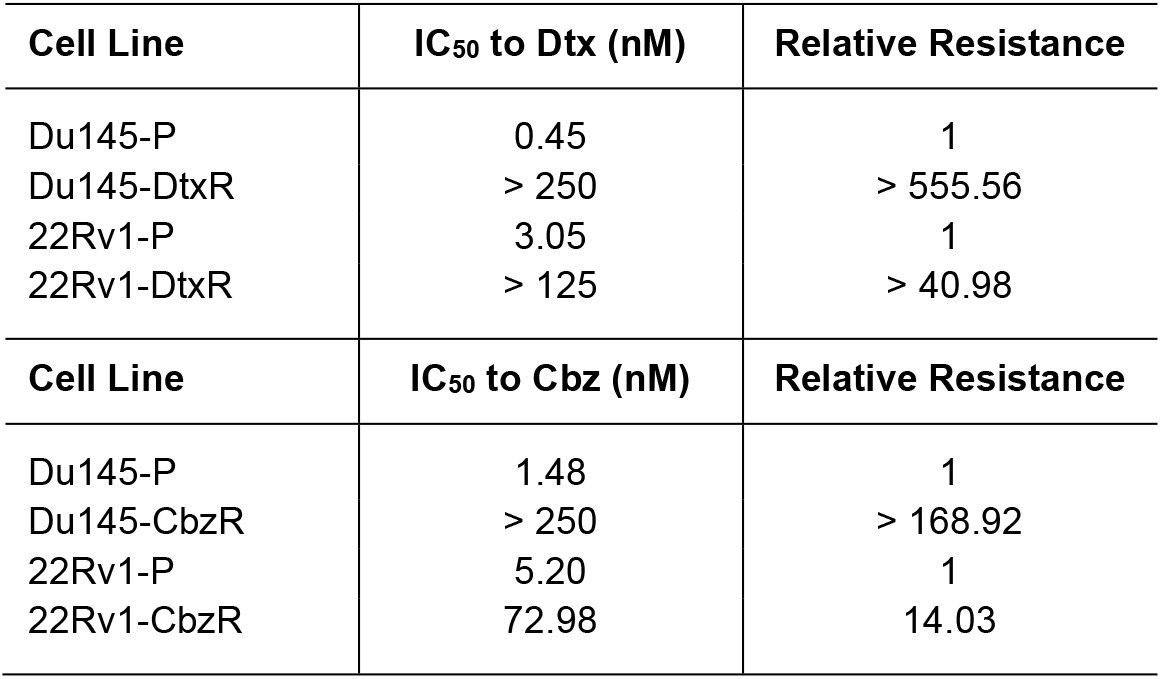
IC_50_ values and relative resistance of Dtx/Cbz resistant Du145 and 22Rv1 cells compared to parental cells. Cells were incubated with increasing concentrations of Dtx/Cbz for 72 h prior to the CTG assay.

| Cell Line | IC<sub>50</sub> to Dtx (nM) | Relative Resistance |
| --- | --- | --- |
| Du145-P | 0.45 | 1 |
| Du145-DtxR | > 250 | > 555.56 |
| 22Rv1-P | 3.05 | 1 |
| 22Rv1-DtxR | > 125 | > 40.98 |
| Cell Line | IC<sub>50</sub> to Cbz (nM) | Relative Resistance |
| Du145-P | 1.48 | 1 |
| Du145-CbzR | > 250 | > 168.92 |
| 22Rv1-P | 5.20 | 1 |
| 22Rv1-CbzR | 72.98 | 14.03 |

**Figure 1.**
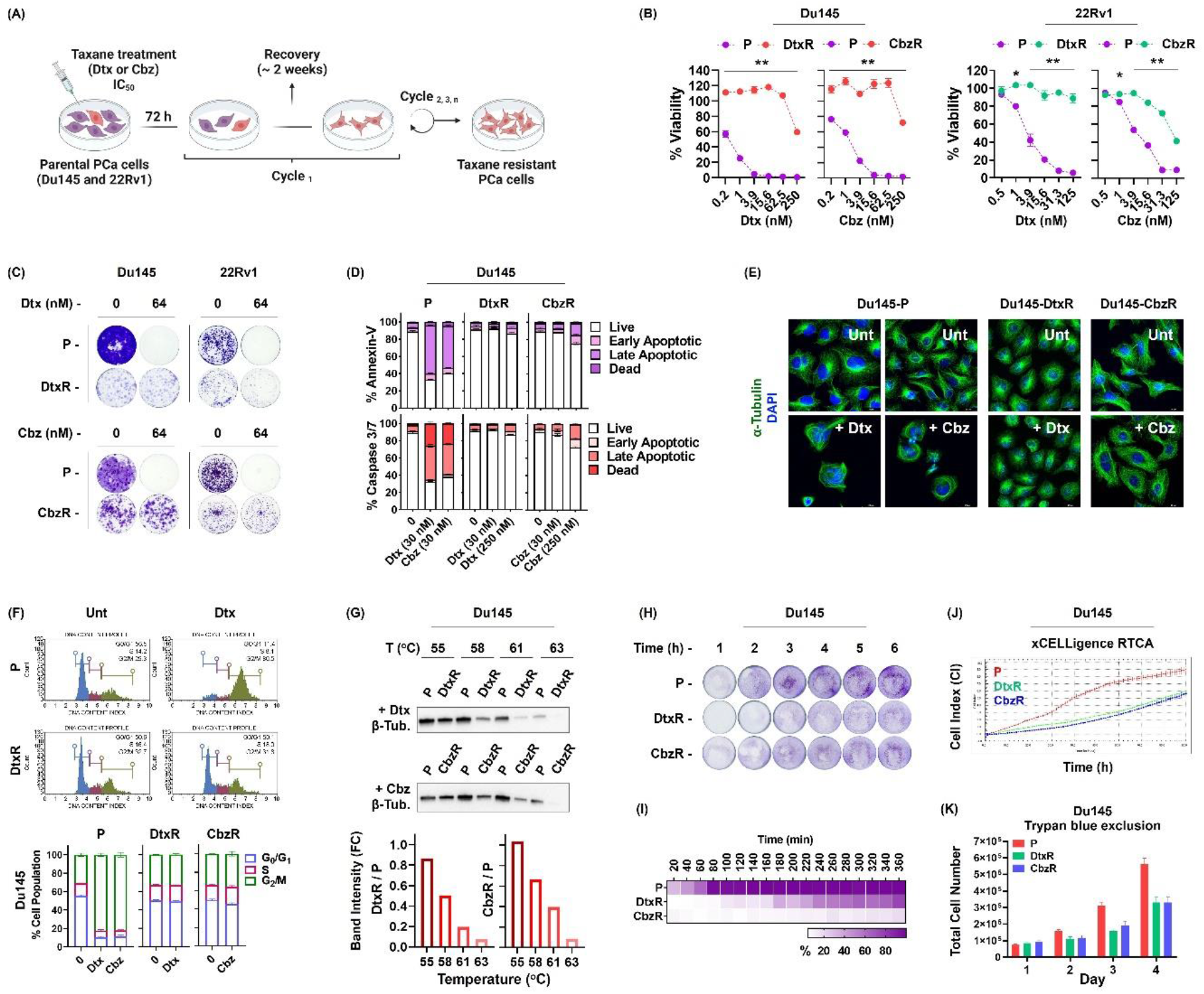
**(A)** Timeline and schema for establishing taxane resistant cell models. Figure was made in BioRender. **(B)** The dose dependent effects of Dtx or Cbz on the cell viability of parental (P) and resistant (R) PCa cell lines. The results were obtained by using CTG assay (72 h) and expressed as mean ± SEM. (*) p < 0.01 and (**) p < 0.0001 indicate significant differences in cell viability between parental and resistant cells. **(C)** Representative clonogenic images were obtained by treating cells with Dtx or Cbz for 72 h and the colony formation ability was analyzed 10-15 days after drug exposure. **(D)** Flow cytometry analysis of cell death in taxane treated Du145-P and -R cells. **(E)** Immunofluorescence staining of α-tubulin (green) in Du145-P and -R cells treated with the indicated taxane (30 nM) for 24 h. Nuclei (blue) were stained with DAPI. **(F)** The cell cycle distribution of Du145-P/R cells treated with Dtx/Cbz (15 nM). Representative DNA content histograms (upper panel) and quantifications are shown (lower panel). **(G)** CETSA for in-cell β-Tubulin target engagement. Western blots showing thermostable β-Tubulin following indicated heat shocks in the presence of Dtx (62.5 nM) or Cbz (31.5 nM) in Du145-P/R cells. CETSA images were quantified by using ImageJ software (shown below). **(H)** Adhesion capacity of Du145-P/R cells. Attached cells were fixed at the indicated time points and were stained with crystal violet. Quantification was performed using ImageJ software **(I)**. **(J)** Real time monitoring of cell growth of Du145-P/R cells with the xCELLigence system. **(K)** The growth rate of Du145-P/R cells was also determined by counting live cells by trypan blue staining. **P:** parental, **R:** resistant.

Having successfully established taxane resistant PCa models, we noted that resistant cells remained adherent, and morphologically heterogeneous, appearing as round, oval or spindle shaped cells that were similar to parental cells (**Sup. Figure 2A**), suggesting that epithelial to mesenchymal transition (EMT) programmes were not triggered. Supportingly, resistant cells exhibited no change in classical EMT markers, while differences between different cell lines existed (**Sup. Figure 2B**). One of our observations with these cells was the differences in their adhesion properties. As shown in **Figure 1H and Sup. Figure 1E** the adhesion of resistant cells required more time compared to parental cells. The other significant difference observed was that resistant cells proliferate more slowly compared to parental cells. We first measured the proliferation rate of parental and resistant cells using a real time analysis with the xCELLigence system (**Figure 1J, Sup. Figure 1F**). In parallel, the proliferation rate of the cells was assessed by trypan blue staining (**Figure 1K, Sup. Figure 1G**). Both methods gave similar results, the growth rate of resistant cells was similar to each other (DtxR vs. CbzR) but differed greatly from the parental cells (TaxR vs. P) as they proliferated at a markedly slower rate.

Interestingly, while the cells were cross-resistant against other chemotherapeutics such as paclitaxel and doxorubicin (**Sup. Figure 3A**), they became more susceptible to platinum group drugs (**Sup. Figure 3B**), suggesting that a common mechanism acted to detoxify drugs that are substrates of drug efflux pumps, at the expense of becoming more vulnerable towards other drugs. All these data have brought us to a point where we can elucidate the mechanism underlying taxane resistance in our well-characterized PCa cell models.

### Transcriptome profiles of taxane-sensitive and -resistant PCa cells

We conducted a whole transcriptome analysis between taxane-sensitive and -resistant Du145 cells to unravel potential molecular mechanisms of resistance. Principal Component Analysis revealed distinct clustering of replicates from each group, demonstrating clear separation based on global transcriptome profiles **(Sup. Figure 4).**

Differential expression analysis of RNA-seq data revealed significant changes in 2955 genes in the Du145-DtxR cells (927 upregulated and 2028 downregulated) and 1251 genes in the Du145-CbzR cells (364 upregulated and 887 downregulated) compared to their sensitive counterparts **(Figure 2A).** Among the differentially expressed genes (DEGs), the top 10 upregulated and downregulated genes in Du145-DtxR Du145-CbzR cells are listed in **Sup. Table 2** and **3**, respectively. The transcriptomic profile was further analyzed using gene set enrichment analysis (GSEA). The ranked gene sets were tested on the hallmark gene sets, which represent well-defined biologic states and processes (MSigDB, Broad Institute). The relative normalized enrichment scores (NES) demonstrated that MYC signaling, unfolded protein response (UPR), E2F and NF-κB signaling were the most significantly enriched gene sets in Du145-DtxR cells **(Figure 2B, C).** The genes demonstrating core enrichment within these gene sets are listed in **Sup. Table 4**.

**Figure 2.**
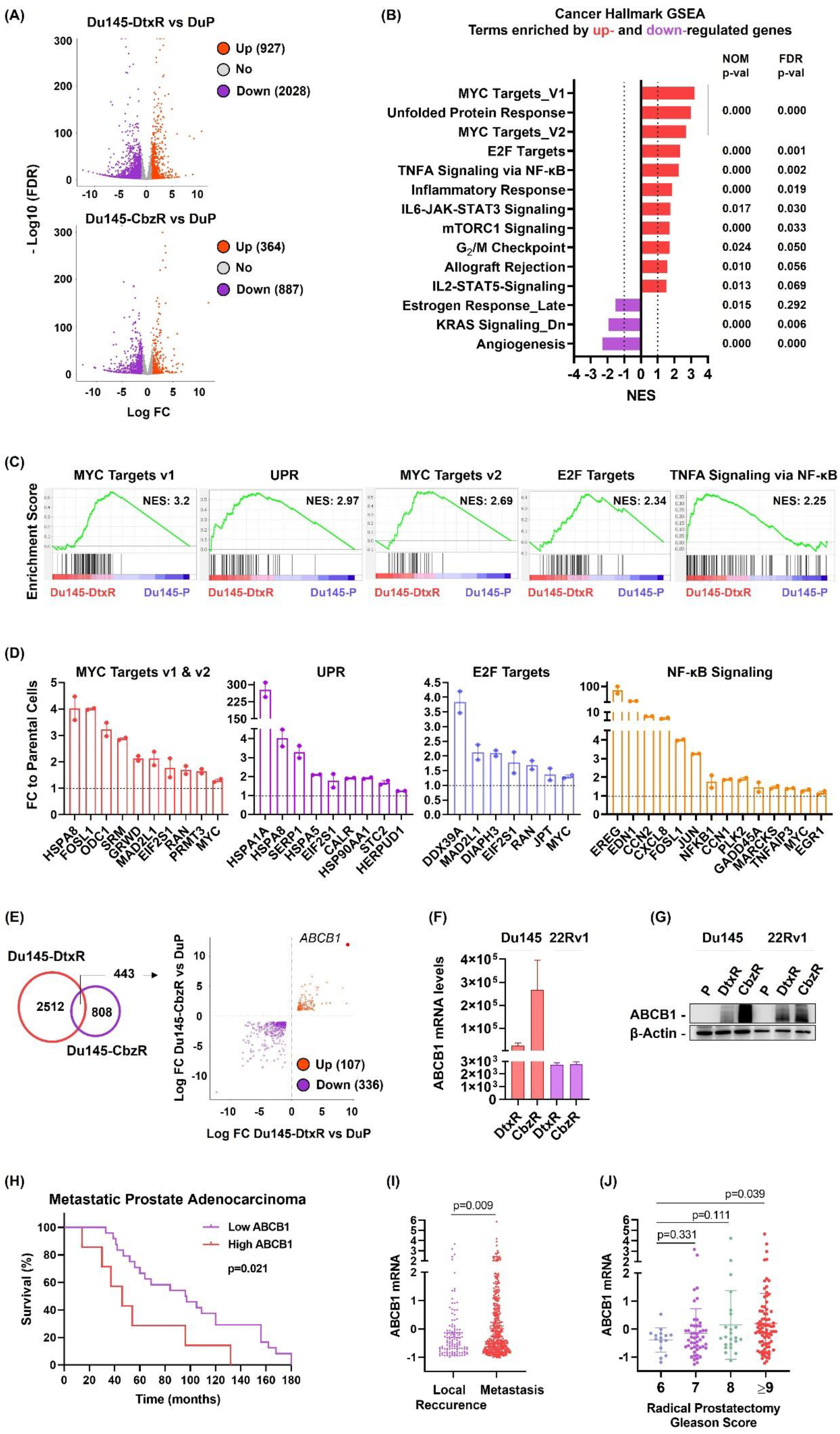
**(A)** Volcano plots showing the DEGs (FDR < 0.05, and Log_2_FC ≥ 1 or ≤ -1) between parental and resistant Du145 cells. The red, purple, and gray scatters indicate upregulated, downregulated and no DEGs between the indicated groups, respectively. **(B)** Gene set enrichment (GSEA) analysis using hallmark gene sets from the MSigDB revealed the upregulation of genes involved in MYC signaling, unfolded protein response (UPR), NF-κB and E2F signaling (FDR < 0.05, and Log_2_FC ≥ 0.5 or ≤ -0.5). GSEA was performed using the h.all.v2023.1.Hs.symbols.gmt dataset in the MsigDB database. **(C)** Enrichment plots for top five gene sets enriched in GSEA Hallmark analysis. **(D)** In-house validation of enriched core genes within the indicated gene sets. **(E)** DEGs (FDR < 0.05, and Log_2_FC ≥ 1 or ≤ -1) from resistant cells were further distributed based on their expression changes in the same direction (up or down). The red and purple scatters indicate up- and down-regulated DEGs, respectively, in resistant Du145 cells. **(F)** Expression levels of ABCB1 mRNA were determined by qRt-PCR. Data is the mean ± SEM. **(G)** ABCB1 protein expression levels in parental and taxane resistant PCa cells were determined by western blotting. **(H)** Comparison of patient survival based on ABCB1 expression levels using a Kaplan-Meier plot (Metastatic Prostate Adenocarcinoma, MCTP). **(I)** ABCB1 mRNA levels in clinical specimens (TCGA-PRAD) were investigated by using cBioPortal and its relationship with the occurrence of local recurrence and metastasis (Pan-cancer Analysis of Advanced and Metastatic Tumors, BCGSC); **(J)** and gleason score (Metastatic Prostate Adenocarcinoma, SU2C/PCF Dream Team) were shown as scatter plots.

GO analysis revealed significant enrichment in biological processes related to ribosome biogenesis, mitochondrial gene expression, and chaperone binding, indicating an increased demand for protein synthesis, efficient mitochondrial function, and proper protein folding in Du145-DtxR cells (**Sup. Figure 5).** Consistently, cellular components such as the chaperone complex, ER protein-containing complex, and mitochondria were found to be enriched, indicating an active cellular state characterized by robust protein synthesis, efficient ribosome assembly, and chaperone-mediated protein quality control mechanisms. Additionally, the negative enrichments indicate alterations in developmental processes, neuronal signaling, calcium regulation, and ECM interactions **(Sup. Figure 5)**.

To confirm the RNA-seq data, another batch of RNA samples were extracted and subjected to qPCR analysis for the validation of core genes within the top 5 gene sets. The selection of these genes for validation was based on two criteria: their ranking in the GSEA gene order **(Sup. Table 4)** and their documented relevance to taxane or drug resistance according to literature searches **(Sup. Table 5).** mRNA expression of selected genes was upregulated in taxane-resistant cells, confirming RNA-seq results **(Figure 2D).**

The enrichments of "MYC Targets_V1" and "E2F Targets" were also observed in Du145-CbzR cells, similar to Dtx-resistant cells. This suggests a potential involvement of MYC and E2F signaling pathways in mediating resistance mechanisms across different taxanes **(Sup. Figure 6A, B).** The top positive enrichment of the "Peroxisome" pathway indicates an elevated activity or upregulation of peroxisomal functions, which have been reported to support cancer cell survival (Cai et al., 2018; Dahabieh et al, 2017). GO analysis revealed positive enrichments in mitotic processes, including spindle organization, sister chromatid segregation, and microtubule cytoskeleton organization, which align with the mechanism of taxanes targeting microtubules to disrupt cell division **(Sup. Figure 7).** The positive enrichments in membrane-related MFs, including lipid and phospholipid transporter activities, ATP hydrolysis, and primary active transmembrane transporter activities, suggest that these activities are upregulated in these cells, possibly due to the altered membrane composition and increased ATP-dependent transmembrane transport requirements associated with taxane resistance. The genes demonstrating core enrichment within these gene sets are shown in **Sup. Figure 6B**, and their expression was further validated **(Sup. Figure 6C)**.

The data clearly indicate the presence of common patterns, including the upregulation of MYC and E2F signaling and the negative enrichment of ECM binding, in both cells. However, there is also some degree of variation in the underlying mechanisms within each cell. Therefore, we utilized a Venn diagram to identify the common gene pool specific to taxane resistance in both cells, categorizing the DEGs based on their expression changes (up or down) (**Figure 2E**). Our analysis revealed that 107 genes were commonly upregulated and 336 were downregulated in taxane resistant cells. Notably, ABCB1 emerged as the gene with the highest transcriptional change in both resistant cells (**Figure 2E**).

### ABCB1 overexpression in taxane resistant PCa cells

The top transcriptional change took place for the ABCB1 gene, with a 10.3 and 12.9-fold (log_2_ scale) upregulation in Du145-DtxR and Du145-CbzR, respectively, compared to parental cells. After validating the transcript expression of ABCB1 **(Figure 2F)**, we confirmed the corresponding protein upregulation in all taxane resistant cells (**Figure 2G)**. The patient data analysis highlighted the clinical significance of ABCB1 expression, with ABCB1-high patients showing unfavorable outcomes in terms of OS **(Figure 2H).** Furthermore, the analysis of advanced and metastatic tumors revealed a significant relationship between higher ABCB1 mRNA levels and an increased incidence of metastasis compared to patients with local recurrence **(Figure 2I).** In another dataset of patients with metastatic disease, an association was observed between higher Gleason scores and increased ABCB1 mRNA expression **(Figure 2J).** Taken together, these findings provide evidence supporting a potential relationship between ABCB1 expression and disease progression.

One of the potential causes for the increased expression of ABCB1 is gene amplification. Therefore, we examined ABCB1 gene copy numbers in all PCa cell lines. ABCB1 amplification was evident only in Du145-CbzR cells (**Sup. Figure 8**). We then assessed the taxane susceptibility in the presence of verapamil (a P-gp inhibitor). Resensitization was observed following combinatorial verapamil treatments (**Sup. Figure 9A).** Similar results were obtained with the third generation ABCB1 inhibitors, elacridar and zosuquidar (**Sup. Figure 9B).** In addition, verapamil also caused a significant reversal of chemoresistance against paclitaxel and doxorubicin (**Sup. Figure 10**). Despite the presence of cross-resistance, we hypothesized that decreased efflux transporter expression may contribute to the observed platinum sensitivities in resistant cells **(Sup. Figure 3)**. RNA-seq analysis of cisplatin efflux transporters ABCA1 (Du145-DtxR Log_2_FC: -2.62, Du145-CbzR Log_2_FC: -1.85) and ABCA7 (Du145-DtxR Log_2_FC: -0.94, Du145-CbzR Log_2_FC: -1.18), followed by validation, revealed a common decrease in ABCA7 expression in all resistant PCa cells, providing evidence supporting this mechanism **(Sup. Figure 11)**.

To assess the importance of ABCB1 expression in taxane resistance, we used siRNA to target ABCB1 in resistant Du145 cells **(Sup. Figure 12A, B)**. However, we encountered difficulties in achieving effective knockdown of ABCB1 using the same siRNA in resistant 22Rv1 cells. Therefore, we proceeded to knock out ABCB1 expression in these cells **(Sup. Figure 12C, D).** The inhibition of ABCB1 expression in all resistant cells resulted in decreased survival at the doses of taxanes that showed minimal cytotoxicity in the previous results, highlighting the significance of ABCB1 expression in mediating taxane resistance **(Sup. Figure 12E-H)**.

The intracellular accumulation of calcein-AM was examined to confirm the functional activity of the ABCB1 in taxane resistant cells. Calcein-AM, a non-fluorescent substrate of ABCB1, is cleaved by intracellular esterases to produce fluorescent (green) calcein. Calcein-AM efflux is increased in the presence of ABCB1, thus limiting the fluorescent calcein production **(Sup. Figure 13A)**. Consistent with ABCB1 expression, majority of the parental Du145 and 22Rv1 cells retained the dye, while very few resistant cells were positive for green fluorescent calcein, indicating its efflux from the cells **(Sup. Figure 13B, minus columns, P vs R)**. Verapamil treatment did not influence calcein-AM efflux in drug-sensitive cells lacking ABCB1 expression **(Figure 2G)** but caused a robust increase in calcein positive cells for all of the resistant Du145 and 22Rv1 cells (**Sup. Figure 13B, - vs + images for each cell line**). Additionally, the cells were treated with epothilone B (Epo B), a tubulin targeting drug which is not a P-gp substrate. Resistant cells exhibited similar viability curves to parental cells upon Epo B exposure, providing further evidence that the observed difference between the resistant and parental cells was predominantly dependent on ABCB1 (**Sup. Figure 14**).

### Episensitization of taxane resistant PCa cells

Given the epigenetic changes that contribute to taxane resistance in PCa, we performed an epigenetic drug library screen to uncover potential targets and vulnerabilities in PCa cells **(Figure 3A)**. The epigenetic drug library has 145 compounds including a majority of currently available agonists and antagonists of epigenetic regulatory enzymes. The top 5 classes of epigenetic targets that induced the most pronounced resensitization when applied to resistant cells were identified as CBP/p300, BRPF, Menin-MLL, PRMT5, and SIRT1 **(Figure 3B, C)**. The efficacy of the epigenetic drugs was further validated using cell viability assays at different concentrations, revealing a restoration in taxane susceptibility with increasing doses of epidrugs **(Sup. Figure 15)**. However, subsequent investigation showed that expressing SIRT1 did not have a significant impact on resensitization of taxane-resistant cells. (**Sup. Figure 16A, B)**. Additionally, silencing CBP, a well-known transcriptional coactivator with histone acetyltransferase activity, resulted in a complete loss of colony-forming ability of cells, indicating their essentiality for PCa cell survival (**Sup. Figure 16C)**. On the other hand, our lab has ongoing studies on Menin-MLL and PRMT5, which are being investigated in detail and will be reported separately. Therefore, the findings from our epigenetic drug library screen led us to prioritize BRPF proteins as a potential target for overcoming taxane resistance in PCa. We also identified a knowledge gap regarding the functions of BRPF epiregulators and the efficacy of BRPF inhibitors **(Figure 3D)** in cancer drug resistance, which drove us to highlight these proteins as promising and novel targetable epiregulators in this context.

**Figure 3.**
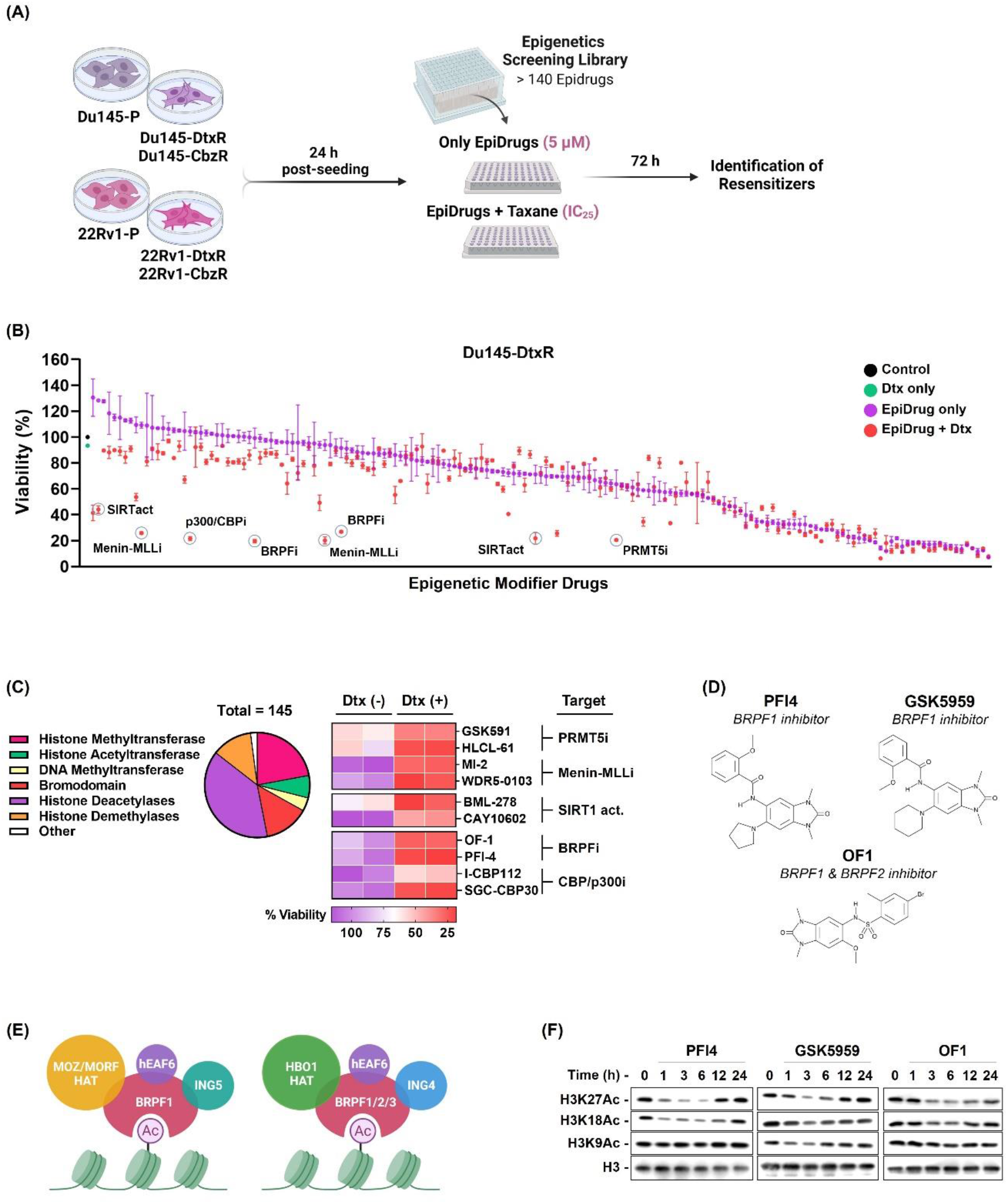
**(A)** Schematic view of the epidrug library screening. The figure was generated using BioRender software. **(B)** Screening of 145 targeted drugs to identify Epigenetic modulators capable of overcoming taxane resistance was performed using the SRB assay and the result obtained in Du145-DtxR cells is shown representatively. Purple dots show the cell viabilities upon the standalone use of each epidrugs, while the red dots indicate the cell-killing effect of their combination with Dtx. Re-sensitizers ranked in the top 5 groups are highlighted in circles. **(C)** Pie Chart illustrates the epigenetic proteins targeted by the small molecules (epidrugs) in the library. Epidrugs were screened for resistance reverter activity which resulted in the identification of 5 classes of epigenetic targets (PRMT5, Menin-MLL, SIRT1, BRPF and CBP/p300) that induced the most pronounced resensitization when applied to resistant cells. **(D)** Molecular structure of BRPF inhibitors. **(E)** Cartoon representation of the BRPF family and **(F)** demonstration of the efficacy of BRPF inhibition on histone acetylation by western blotting.

### Targeting BRPF epiregulators in taxane resistant PCa cells

The BRPF epigenetic reader family (BRPF1, BRPF2, and BRPF3) acts as scaffolds for assembling HAT complexes of MOZ/MORF and HBO1 families with ING5 and Eaf6, carrying these to chromatin via its bromodomain (Meier et al., 2017) **(Figure 3E).** Both BRPF2 and BRPF3, despite their high sequence similarity to BRPF1, have a preference for forming complexes with HBO1 instead of MOZ and MORF, suggesting a distinct functional behavior of BRPF1. Moreover, BRPF1 directs histone acetylation towards H3 instead of H4, indicating its pivotal role in controlling the substrate specificity of HBO1 (Lalonde et al., 2013). BRPF1 is known to be indispensable for embryonic development; however, its involvement in cancer, with the exception of limited studies, remains poorly characterized.

We validated these hits by screening each BRPF inhibitor in a wider dose range and confirmed the resensitizing effect as resistant cells became more responsive to taxanes with increasing doses of these inhibitors **(Figure 4A** and **Sup. Figure 17A)**. Additionally, the observed effect was found to be synergistic, as evident by combination index (CI) values **(Figure 4B)**. These findings were further supported by colony formation assays **(Figure 4C)**. Interestingly, parental cells did not show any increased sensitivity to epidrug treatments indicating that BRPF inhibitors have the potential to selectively target resistant cells **(Sup. Figure 17B).** To confirm whether the growth inhibition observed in response to the drug combinations was due to cell death, we assessed apoptotic markers. Combining taxanes and epidrugs resulted in increased annexin-V staining and caspase 3/7 activation compared to using either drug alone **(Figure 4D).** Furthermore, the G_2_/M arrest of resistant PCa cells by BRPF inhibitors provides further support for the restoration of taxane sensitivity **(Figure 4E** and **Sup. Figure 17C).** Lastly, increased BRPF1 expression was found to be associated with elevated expression of the cell growth indicator, mKI67, in patients with metastatic PCa, providing evidence for its role in promoting cellular proliferation pathways **(Figure 4F).**

**Figure 4.**
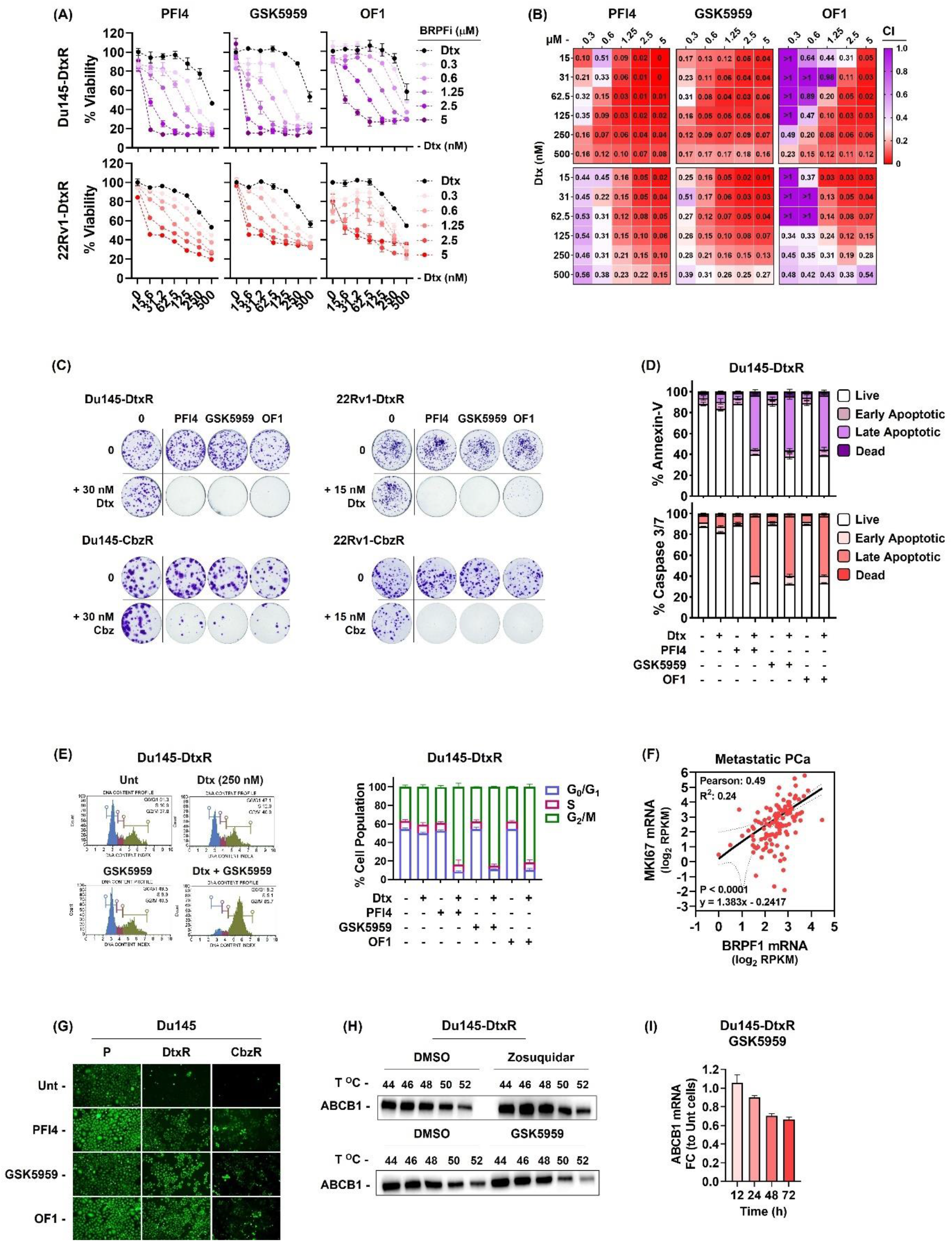
**(A)** Validation dose-response curves of BRPF inhibitors (PFI4, GSK5959 and OF1) on Dtx-resistant cells. Cells were co-treated with indicated drugs (Dtx; 1-250 nM and BRPF inhibitors; 1.25-5 µM) and the results were obtained by SRB viability assay (72 h). The data is expressed as mean ± SEM. **(B)** Heat map representation of the Combination Index (CI) values, with red color indicating a synergistic effect. CI was calculated using the CalcuSyn software. **(C)** Clonogenic images were obtained by treating cells with indicated drugs for 72 h and the colony formation ability was analyzed 10-15 days after drug exposure. **(D)** Flow cytometry analysis of cell death (48 h) and **(E)** cell cycle distribution (24h) in resensitized Du145-DtxR cells. **(F)** The expression of BRPF1 showed a positive correlation with MKi67 expression in metastatic PCa. **(G)** Calcein retention assay was performed in the absence or presence of BRPF inhibitors (5 µM, 24h). **(H)** CETSA for in-cell ABCB1 engagement. Western blots showing thermostable ABCB1 following indicated heat shocks (44°C, 46°C, 48°C, 50°C and 52°C) in the presence of Zosuquidar (5 µM) and GSK5959 (5 µM) in Du145-DtxR cells. **(I)** The expression of ABCB1 was evaluated by qRt-PCR at the indicated time points following treatment with GSK5959 (5 µM). **P:** parental, **R:** resistant.

Based on the apparent role of ABCB1 in taxane response (**Sup. Figure 11)**, it is reasonable to speculate that these inhibitors could interfere with its function. To test this, we performed calcein efflux assay with BRPF inhibitors. Despite reduced calcein accumulation in resistant cells due to high ABCB1 expression, green fluorescence significantly increased with all BRPF inhibitors, supporting their potential role in inhibiting ABCB1 function **(Figure 4G, Sup. Figure 17D)**. It could be assumed that these epidrugs can be effective by binding to ABCB1. To investigate the potential binding of BRPF inhibitors with ABCB1, we performed CETSA against ABCB1. Zosuquidar was used as a positive control and not surprisingly, it showed an ABCB1-binding compared to DMSO **(Figure 4H, Sup. Figure 17E)**. On the other hand, BRPF inhibitors did not cause any thermal shift, therefore we concluded as they do not engage with ABCB1 physically. The restoration of taxane sensitivity in resistant cells might be attributed to downstream mechanisms rather than direct binding to ABCB1. Indeed, a time-dependent decrease in the expression of ABCB1 was observed upon treatment with GSK5959 **(Figure 4I).** Additionally, the lack of cytotoxic effects observed in RPE-1 cells upon treatment with BRPF inhibitors provides promising evidence for their potential therapeutic application **(Sup. Figure 17F).** The differential response to epidrug treatments between sensitive and resistant cells can be attributed to the expression of ABCB1. Indeed, knocking out ABCB1 in the resistant cells abolished the effectiveness of BRPF inhibitors, suggesting that the effect of BRPF inhibitors is dependent on the presence of ABCB1 **(Sup. Figure 18)**.

In order to investigate whether the effect of BRPFi can be recapitulated using gene targeting, we employed both CRISPR guided knockout and siRNA mediated silencing strategies. Several attempts to generate BRPF knock out cells failed, as resistant cells did not survive following introduction of targeted gRNAs indicating that the activity of BRPF genes became essential for the resistance phenotype. On the other hand, short term depletion, as in the case of RNA interference, was applicable. The colony formation assay showed that knockdown of BRPF1 suppressed the ability to form colonies **(Sup. Figure 19A).** However, this effect was observed to be more intense in resistant cells, once again showing the essentiality of BRPF1 in resistant cells. Indeed, stable and specific knock-down of BRPF1 led to a higher number of apoptotic cells in Dtx-resistant cells **(Sup. Figure 19B, C).**

While not as pronounced as the effect seen with pharmacological inhibition, knockdown of BRPF1 also led to the resensitization of cells to taxanes **(Figure 5A, B).** We also noted a decrease in ABCB1 expression upon BRPF1 knockdown **(Figure 5C).** The observed decrease in ABCB1 expression with both pharmacological and genomic suppression implies a potential role of BRPF1 in transcriptional regulation. In order to test whether BRPF1 directly regulates ABCB1 expression, we analyzed the enrichment of BRPF1 on the promoter of ABCB1 in Du145-DtxR cells using ChIP assay. We found that, in comparison with IgG and gene desert region controls, BRPF1 was indeed enriched at the promoter region of ABCB1 **(Figure 5D).**

**Figure 5.**
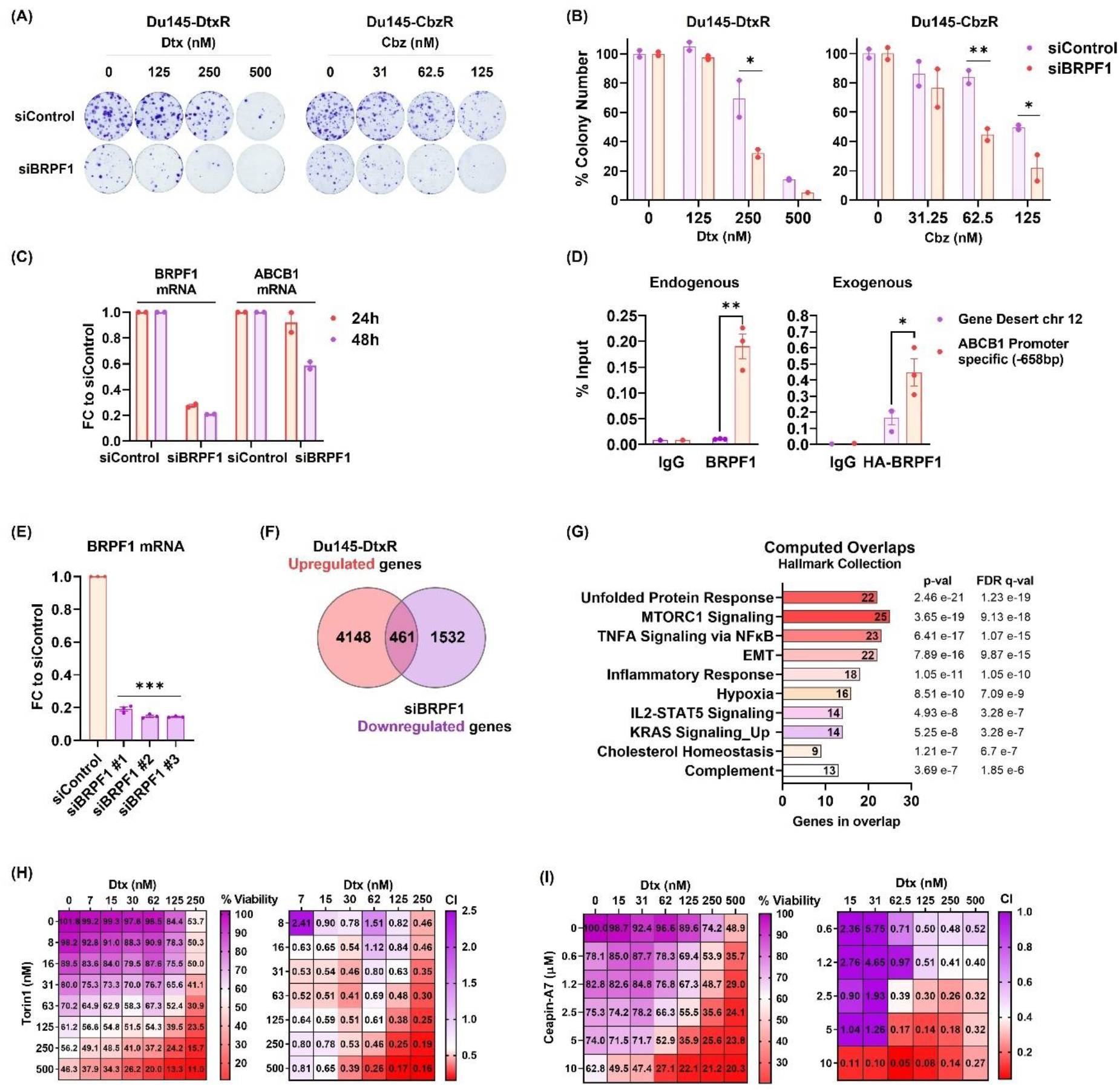
**(A)** Taxane response of siBRPF1 treated resistant cells evaluated by colony formation assay. Representative images and **(B)** quantifications are shown. **(C)** The expression of ABCB1 was evaluated by qRt-PCR at the indicated time points following treatment with siBRPF1 in Du145-DtxR cells. **(D)** ChIP-qPCR showing BRPF1 enrichment at the *ABCB1* promoter in Du145-DtxR cells expressing endogenous BRPF1 (*left panel)* and exogenous HA-tagged BRPF1 (*right panel*). BRPF1 enrichment at a control region (*Chr12 gene desert*) is also shown. Data are shown as percentage of ChIP input; dots represent individual biological replicates, bars represent mean replicates. **(E)** BRPF1 mRNA levels of RNA-seq samples from siControl and siBRPF1 in Du145-DtxR cells. **(F)** Venn diagram showing the number of genes (intersection, 461) whose expression decreased after silencing of BRPF1 among genes with increased expression in Du145-DtxR cells (vs Du145-P). **(G)** Computed overlaps of the 461 genes in the Hallmark Collection of GSEA (MSigDB) database. **(H)** The efficacy of Torin1 (mTORC1/2 inhibitor, 8-500 nM) on Du145-DtxR cells was determined by CTG assay and represented as a heat map. **(I)** The efficacy of Ceapin-A7 (ATF6α inhibitor, 0.6-10 µM) on Du145-DtxR cells was determined by SRB assay and represented as a heat map. The Combination Index (CI) was calculated using the CalcuSyn program. Statistical significance denoted as (*) p < 0.05, (**) p < 0.01, and (***) p < 0.0001.

To reveal a BRPF1 related mechanism, we wanted to examine the transcriptome of siBRPF1 treated Dtx-resistant cells **(Figure 5E).** We limited the gene pool of interest by filtering the upregulated genes whose expression was previously determined in resistant cells (relative to parental cells) **(Figure 2A)** but showed an opposite direction of regulation after siBRPF1 treatment **(Figure 5F** and **Sup. Figure 20).** Among the pathways/signal transductions in which a total of 461 genes overlap in the GSEA (MSigDB) database (Hallmark Collection), mTORC1 and UPR have emerged as the most significant overlaps **(Figure 5G, Sup. Table 6)**. Torin1 (an inhibitor of mTORC1/2) and Ceapin-A7 (an inhibitor of ATF6α) were used to assess the essentiality of these pathways in reversing resistance. Administration of the inhibitors restored the sensitivity of resistant cells to Dtx, while having no effect on the response of sensitive cells, mimicking the effect observed with BRPF inhibitors **(Figure 5H, I,** and **Sup. Figure 21)**. These observations suggest that BRPF1 may exert a regulatory role in modulating ABCB1 expression through its interaction with the UPR and mTORC1 signaling pathways. Indeed, the combination of ABCB1 silencing and mTOR inhibition enhances the sensitization of cells to taxanes, suggesting a mechanism involving multiple downstream pathways regulated by BRPF1 **(Sup. Figure 22)**. These results indicate the involvement of diverse mechanisms in overcoming taxane resistance. The combined targeting of ABCB1 and mTOR pathways may act through complementary mechanisms, such as inhibiting drug efflux and modulating intracellular signaling cascades, ultimately leading to the restoration of drug sensitivity.

Upon examining the functional clusters of upregulated genes with siBRPF1 treatment, it was reasonable to observe the emergence of gene sets related to PCa, such as "androgen response” or “mitotic spindle” and “G_2_M checkpoint", given the mechanism of action of taxanes **(Sup. Figure 20).** Comprehensive studies are needed to investigate the significance of these genes in taxane resistance **(Sup. Table 7).**

We analyzed publicly available clinical data and initially performed a pan-cancer analysis. We divided the patients into low and high BRPF1 expression groups using the median values determined by the cBioPortal (**Figure 6)**. Our findings demonstrated a significant decrease in the overall survival of patients exhibiting high levels of BRPF1 expression **(Figure 6A)**. Further analysis of the specific cancer types revealed higher BRPF1 expression in lymphoma, while lower expression levels were observed in hepatobiliary, pancreatic, and prostate cancers **(Figure 6B)**. In the analysis of sample types, we observed a higher expression of BRPF1 in local recurrence and metastasis **(Figure 6C)**. These findings suggest that despite being categorized in the low BRPF1 group, the expression of BRPF1 in PCa may be implicated in disease progression. Indeed, we found a significant association between BRPF1 expression levels and PCa progression, with higher expression levels being correlated with increasing Gleason score and tumor stage (**Figure 6F).** Moreover, BRPF1 expression might be a potential recurrence indicator, as higher expression was observed in patients upon recurrence (**Figure 6G).** Supportingly, its expression was found to be elevated in metastatic samples (**Figure 6G).** BRPF1 expression also seems to reflect therapy outcome, as patients with higher expression levels tend to have a poorer response to treatment and more progressive disease (**Figure 6H).** Additionally, it is noteworthy that an increase in BRPF1 expression was observed in patients who were exposed to taxane **(Figure 6H)**, further emphasizing its potential relevance in therapy outcome and disease progression. Overall, these data highlight the clinical significance of BRPF1 expression as a potential biomarker for PCa progression.

**Figure 6.**
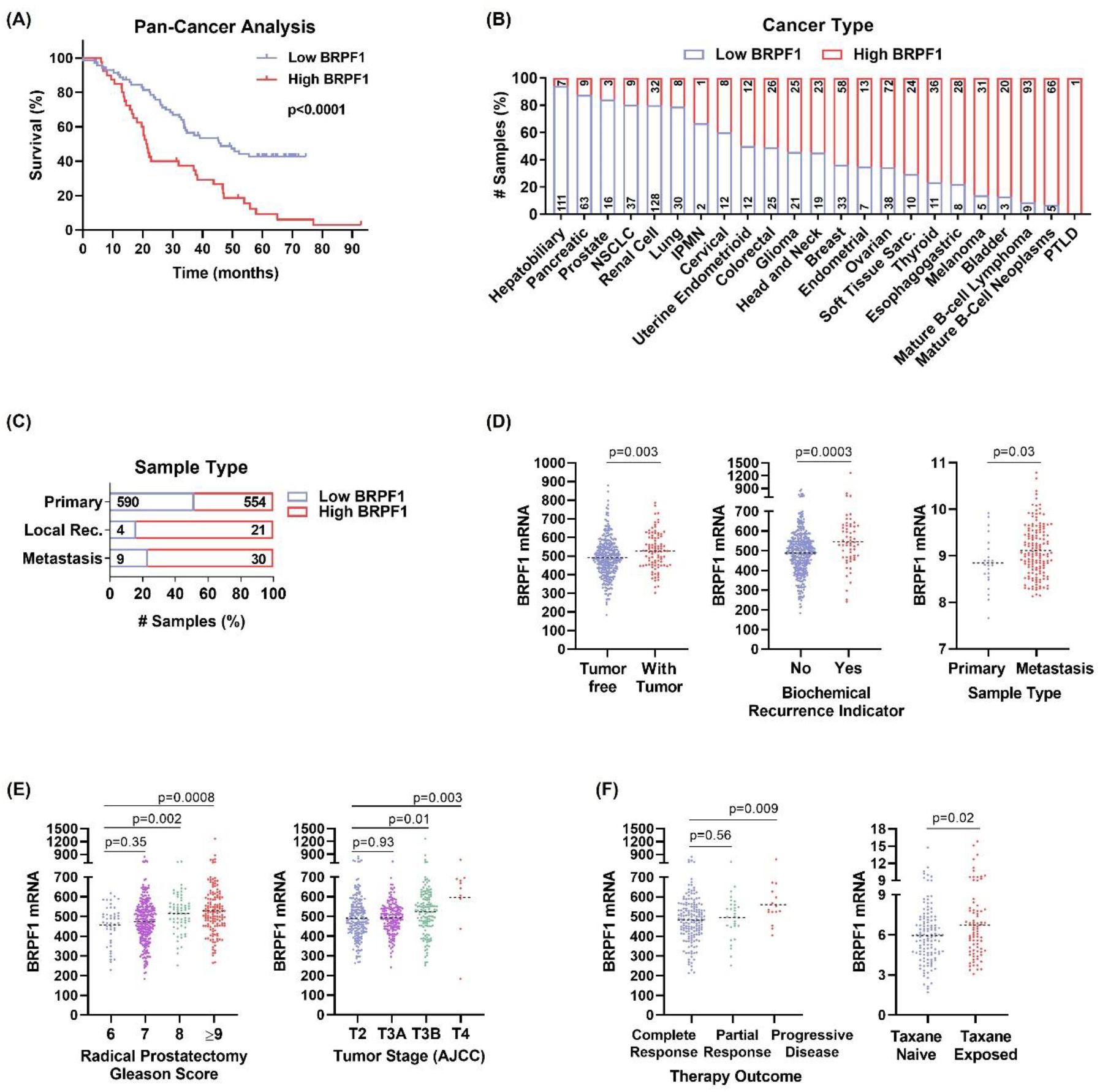
**(A)** BRPF1 mRNA levels were analyzed in the Pan-Cancer Analysis dataset (ICGC/TCGA, Nature 2020) from cBioPortal. Kaplan-Meier plot showing the comparison of patient survival based on BRPF1 expression levels. The patients were classified into Low BRPF1 and High BRPF1 groups based on the expression median. The distribution of BRPF1 expression **(B)** across different cancers and **(C)** sample types. The numbers within the column bars correspond to the sample size. **(D)** BRPF1 mRNA levels in clinical PCa specimens were examined for their correlation with neoplasm status (Prostate Adenocarcinoma, TCGA, PanCancer Atlas), recurrence (Prostate Adenocarcinoma, TCGA, Firehose Legacy), sample type (Prostate Adenocarcinoma, Fred Hutchinson CRC, Nat Med 2016) **(E)** Gleason score (Prostate Adenocarcinoma, TCGA, Firehose Legacy), tumor stage (Prostate Adenocarcinoma, TCGA, Firehose Legacy), **(F)** therapy outcome (Prostate Adenocarcinoma, TCGA, Firehose Legacy), and taxane exposure status (Metastatic Prostate Adenocarcinoma, SU2C/PCF Dream Team, PNAS 2019). The results were presented as scatter plots.

The analysis of BRPF gene expression in our isogenic cells indicated minimal changes overall, with variations observed among PCa cells and taxanes **(Sup. Figure 23A)**. The lack of observed increase in BRPF1 expression in our cells, unlike in patient tissues, could potentially be attributed to the developed drug resistance that has become chronic in cells. Nevertheless, there could be differences in BRPF activities between resistant and parental cells, which warrants further investigation. If this was indeed associated with a chronic state, our hypothesis was that PCa parental cells may acutely increase BRPF expression following taxane exposure, as supported by the observation of BRPF1 upregulation in PCa patients **(Figure 6F)**. Indeed, the treatment with Dtx resulted in a significant increase in the expression of BRPF1 **(Sup. Figure 23B)**. In addition, we also observed an upregulation of ABCB1 expression following Dtx treatment **(Sup. Figure 23B)**.

Our findings provide the first evidence for i) the potential of BRPF inhibitors to overcome taxane resistance and ii) the regulatory role of BRPF1 through direct modulation of ABCB1 expression at the promoter level. Additionally, we showed that BRPF1 may indirectly regulate ABCB1 expression by interfering with UPR and mTORC1 signaling pathways. The graphical abstract summarizing the study is depicted in **Figure 7**.

**Figure 7.**
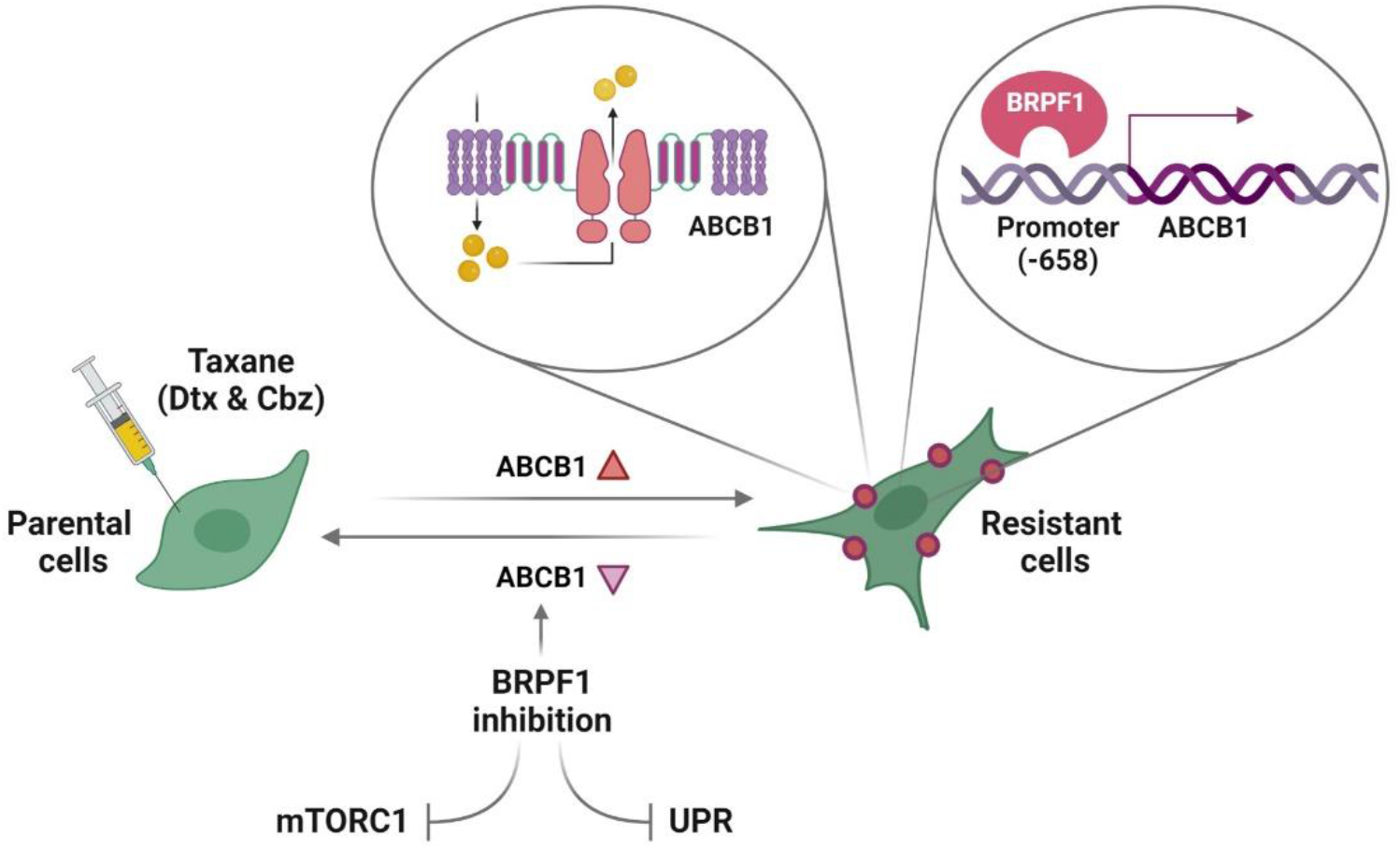
The figure illustrates the major findings of the study.

## Discussion

In this study, we searched for novel epigenetic targets for taxane-resistant CR-PCa treatment. A set of Dtx and Cbz resistant cell lines were derived by pulse selection **(Figure 1A)**. The cells became less responsive to taxanes, exhibited increased IC_50_ values **(Table 1)** and survival **(Figure 1B, C)** and less apoptosis **(Figure 1D)**. Interestingly, while the drug-resistant cells had weaker adhesion potentially conferring advantageous benefits for spreading in the microenvironment and/or metastasis **(Figure 1H, I),** they exhibited relatively slower growth **(Figure 1J, K)** compared to the parental cells which may be due to cell cycle alterations. Not surprisingly, we observed less MT engagement of taxanes in drug-resistant cells **(Figure 1G),** which can be a result of; i. acquired mechanisms that prevent drug accumulation and thus prevent target engagement or ii. tubulin mutations. To elucidate the possible mechanisms falling into the first group, we performed RNA-seq analysis **(Figure 2A-D)** and detected ABCB1 upregulation in all taxane resistant cells **(Figure 2E-G)**. Inhibition of ABCB1 function with small molecules **(Sup. Figure 9-10)** or its expression by using siRNA or gRNA resensitized the resistant cells **(Sup. Figure 12)**. Drug efflux activity of resistant cells was also confirmed by imaging calcein accumulation in the presence of verapamil **(Sup. Figure 13).** Furthermore, by using Epothilone B we showed that the observed difference between the resistant and parental cell lines was ABCB1-dependent **(Sup. Figure 14).** The upregulation of ABCB1 emerges as a frequent and common mechanism underlying Dtx or Cbz resistance in PCa, as highlighted by previous studies characterizing taxane-resistant PCa cells generated through a similar approach to ours (Cajigas-Du et al., 2018, Machioka et al., 2018, Lombard et al., 2019, Seo et al., 2020, Lombard et al, 2021, Linke et al., 2022). Only one of the resistant lines (Du145-CbzR) exhibited copy number amplification **(Sup. Figure 8)**. Lombard et al. demonstrated the activation of the ABCB1 amplicon in taxane-resistant PCa cells and proposed its involvement in resistance beyond ABCB1 alone (Lombard et al., 2021). Their RNA-seq analysis revealed coordinated overexpression of ABCB1-amplicon genes. Among them, targeting the highly overexpressed gene RUNDC3B, in addition to ABCB1, resulted in reduced cell viability and enhanced sensitivity of cells to taxane. This study, along with others as reviewed by Genovese et al., has demonstrated that there is an overexpression or amplification of the genomic region (7q21.12) in taxane-resistant cells (Genovese et al., 2017). Our analysis of the RNA-seq data revealed significant upregulation in 5 out of 11 genes within the ABCB1 amplicon in Du145-CbzR cells, including HNRNPA1P9 (Log_2_FC: 6.7), ABCB4 (Log_2_FC: 5.9), CROT (Log_2_FC: 2.6), TP53TG1 (Log_2_FC: 2.5), and TMEM243 (Log_2_FC: 2.49). Some of these genes were also among the top 10 upregulated genes in Cbz-resistant cells, as shown in **Supp. Table 3**. However, in Du145-DtxR cells, only 2 of the 11 genes were upregulated:HNRNPA1P9 (Log_2_FC: 3.33) and RUNDC3B (Log_2_FC: 1.18), indicating the presence of mechanisms other than gene dosage, such as epigenetic events, in driving the overexpression of ABCB1.

Although several studies have demonstrated the molecular profiling of taxane resistance (Vidal et al., 2015; Lombard et al., 2017; Hongo et al., 2018; Sekino et al., 2019; Jiménez et al., 2020; Hishida et al., 2021), epigenetic modifiers have not been elucidated in taxane resistant CR-PCa. In an attempt to uncover the epigenetic modifiers **(Figure 3A, B)** an epi-drug screen was performed and 5 major classes of targeted molecules (CBP/p300, Menin-MLL, SIRT, PRMT5 and BRPF) appeared as hits with reversion capacity **(Figure 3C).** To our knowledge, this study is the first to report these enzymes as targets and the potential use of their inhibitors in taxane resistant cells **(Figure 3D** and **Sup. Figure 15)**. Targeting BRPFs in resistant cells resulted in resensitization, as evident by CI values of synergy and cell death analyses **(Figure 4A-D)**. Indeed, cells were able to enter G_2_/M arrest upon BRPF inhibition **(Figure 4E)**. These findings showed that, in ABCB1 upregulated cells, taxanes could act in a cellular environment where BRPFs were inhibited and therefore highlight an important role for BRPFs as determinants of taxane susceptibility. Previous studies indicated a role for BRPF1 in bone maintenance (Meier et al., 2017), development of vertebrates (Laue et al., 2008), mouse embryo (You et al., 2015a), forebrain (You et al., 2015b) and fetal hematopoietic stem cells (You et al., 2016), as well as in learning and memory (Xian et al., 2021) and causing neurodevelopmental disorders in humans when mutated (Yan et al, 2017). Moreover, somatic BRPF1 mutations have been identified in sonic hedgehog medulloblastoma (Kool et al., 2014) and pediatric cancers (Huether et al., 2014). Although there are not many studies on the function of BRPFs in cancer, its significance has been noted in liver cancer (Cheng et al., 2021), lower-grade gliomas (Xia et al., 2021) and PCa (Lin et al., 2022). Gene ablation or pharmacological inactivation of BRPF1 significantly attenuated HCC cell growth *in vitro* and *in vivo* (Cheng et al., 2021). Xia et al. identified BRPF1 as a potential drug target in lower-grade gliomas, as inhibiting BRPF1 function or silencing BRPF1 was found to reduce glioma cell proliferation and colony formation (Xia et al., 2021). Lin et al. showed the USP35/BRPF1 axis promoted malignant features of PCa by activating the mevalonate pathway (Lin et al., 2022). Although the oncogenic role of BRPF1 was clearly demonstrated in these studies, its relationship with drug resistance has not been questioned. Therefore, our study is the first to locate BRPFs in cancer drug resistance.

In order to phenocopy BRPF inhibition and rule out potential off-target activities, we depleted BRPF1 and BRPF2 by using RNAi. Based on the very few colonies growing after targeting BRPF2, we were able to test the taxane sensitivity only by targeting BRPF1 **(Sup. Figure 19A)**. Although it was not as potent as we have seen with inhibitors, downregulation of BRPF1 partially reversed taxane resistance **(Figure 5A, B).** One of the explanations for this is that inhibitors may have a wider spectrum of action. While siRNA molecules are effective in decreasing the transcript level, they cannot affect protein levels that are already present in the cellular environment. In addition, the efficacy of multiple targets/signaling pathways that the drug has the potential to affect may not be achieved by targeting a single gene. Unfortunately, despite our several attempts, we failed to generate BRPF1 knockout resistant cells as BRPF1 was very essential for the resistant phenotype. Another plausible explanation is drug engagement to ABCB1, inhibiting its efflux capacity. In the study of Barghout et al., it was demonstrated that various epigenetic probes, including the BRPF1 inhibitor PFI4, potentiate TAK-243 (a ubiquitin-activating enzyme inhibitor) cytotoxicity through off-target ABCG2 inhibition (Barghout et al., 2022). As anticipated, this potentiation did not result in significant alterations in the mode-of-action (i.e., ubiquitylation pathways) or expression levels of ABCG2. However, our study diverges in certain aspects and provides data contrary to the off-target modulation. First, the expression of ABCG2 was downregulated in Du145-CbzR (Log_2_FC: -2.03) cells according to our RNA-seq results, while it was not detected as a DEG in Du145-DtxR cells. Furthermore, CETSA analysis clearly showed that the BRPF inhibitors did not exhibit binding to the ABCB1 protein **(Figure 4H)**. Second, they were able to downregulate ABCB1 mRNA, indicating an interference at the transcriptional level **(Figure 4I)**. Third and most importantly, with the hint of transcriptional regulation, we screened several regions of the ABCB1 promoter and indeed showed that both endogenous and exogenous BRPF1 were enriched at the ABCB1 promoter (-658 bp) **(Figure 5E)** suggesting direct regulation of ABCB1 via BRPF1. Therefore, the inhibition of BRPF, leading to a decrease in ABCB1 levels appears to play a crucial role in reversing taxane resistance in these cells, contrary to an off-target effect.

In order to evaluate the transcriptome changes induced by BRPF1 downregulation and understand how this may revert the resistance phenotype, an RNA-seq analysis was performed. We seeked to find pathways that were differentially regulated in resistant cells and were reverted to more parental-like expression levels upon interference with BRPF1. Among the various pathways that fulfilled these criteria, we observed mTORC1 and UPR signaling pathways as the two most significant pathways showing alterations **(Figure 5H)**. We tested the essentiality of these pathways by using inhibitors. Both mTORC1 and UPR inhibition resensitized resistant cells to taxane and showed no activity on parental cells similar to BRPF inhibition **(Figure 5I, J** and **Sup. Figure 21).** mTOR signaling has been demonstrated to contribute to drug resistance, and the efficacy of its inhibitors has been investigated in clinical trials for PCa, including patients with advanced or resistant disease. Several studies, compiled by Avril et al., have demonstrated the significance of UPR in cancer chemotherapy resistance, while Bonsignore et al. highlighted its potential as a promising druggable target in cancer treatment (Avril et al., 2017; Bonsignore et al., 2023). Further investigation is needed to test the contribution of the genes (listed in **Sup. Tables 6** and **7**) to drug resistance, considering their modulation by BRPF inhibition in these signaling pathways.

Through the examination of clinical data obtained from a pan-cancer study, we categorized patients into low and high BRPF1 expression groups, which revealed a significant association between elevated BRPF1 expression and reduced OS **(Figure 6A).** While it was surprising to find lower BRPF1 expression in PCa among the analyzed cancer types, the elevated expression of BRPF1 in cases of local recurrence and metastasis suggests a potential contribution of BRPF1 to disease progression **(Figure 6B, C).** Indeed, we observed a significant relationship between BRPF1 expression and PCa progression, as higher expression was associated with increased Gleason score and tumor stage, indicating its prognostic potential **(Figure 6E).** Elevated BRPF1 expression was also observed in recurrent and metastatic cases **(Figure 6D)**, suggesting its role as a potential recurrence indicator. Patients with higher BRPF1 expression levels exhibit poor treatment response and more progressive disease **(Figure 6F)**, emphasizing its involvement in therapy outcome and disease progression. Intriguingly, an increase in BRPF1 expression was observed in patients exposed to taxane **(Figure 6F)**. Although we did not observe a significant upregulation of BRPF1 in our taxane-resistant cells **(Sup. Figure 23A)**, we hypothesize that parental cells have the potential to acutely upregulate BRPF expression in response to taxane exposure. Treatment with Dtx indeed led to a significant increase in BRPF1 expression in parental cells **(Figure 23B)**. In addition to the increase in BRPF1, we also observed an upregulation of ABCB1 expression following Dtx treatment. This suggests a potential interplay between BRPF1 and ABCB1, where the upregulation of BRPF1 may contribute to the subsequent upregulation of ABCB1, possibly through transcriptional regulatory mechanisms. In our study, we have provided evidence for the binding of BRPF1 to the ABCB1 promoter, supporting this proposed interaction. Further investigation is needed to elucidate the exact molecular signalings involved in this coordinated response.

## Conclusion

Our results showed that BRPF inhibition could serve as a promising strategy in taxane resistant CR-PCa, and the mechanism of resensitization appears to involve the inhibition of drug efflux in cells that overexpress ABCB1. Since chemical inhibition of ABCB1 through small molecules is not feasible for cancer therapy due to high toxicity, revealing its regulators offers a good option for reducing its activity. In this context, our study herein is the first to identify BRPF1 as an ABCB1 regulator and uncover several epiregulators with the potential to reverse taxane resistance in CR-PCa.

## Acknowledgement

The authors gratefully acknowledge the use of the services and facilities of the Koç University Research Center for Translational Medicine (KUTTAM), funded by the Presidency of Turkey, Presidency of Strategy and Budget. The following figures were created using BioRender.com and licensed for publication (Figure 1: FK25PK7QVH, Sup. Figure 1: RR25PKC49Z, Sup. Figure 13: PA25PKCI9L, Figure 3A: FP25PKCQ80, Figure 3E: QP25PKCYLM, Figure 7: DM25PKDVVA).

---

**Sup. Table 1.**
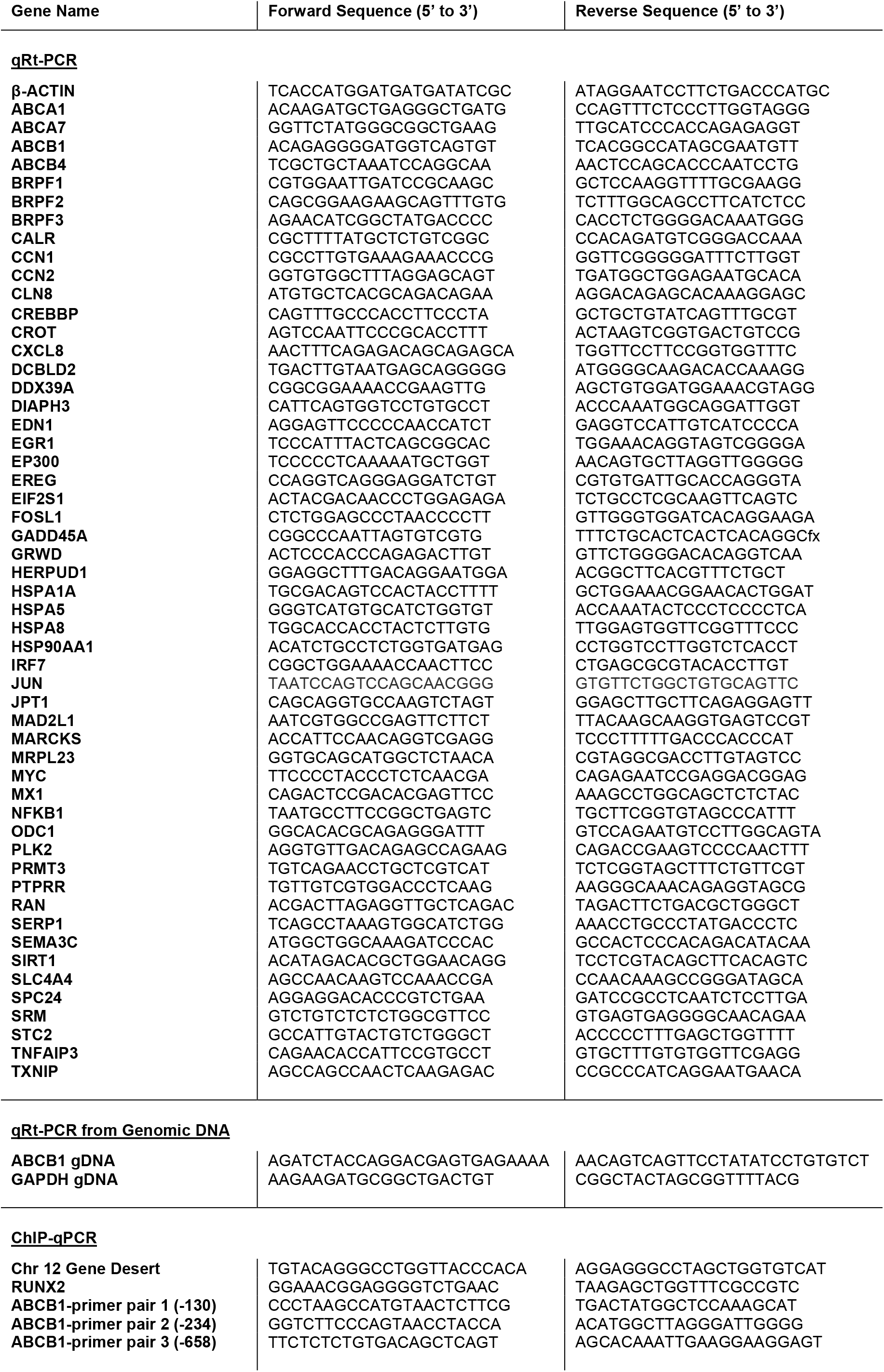
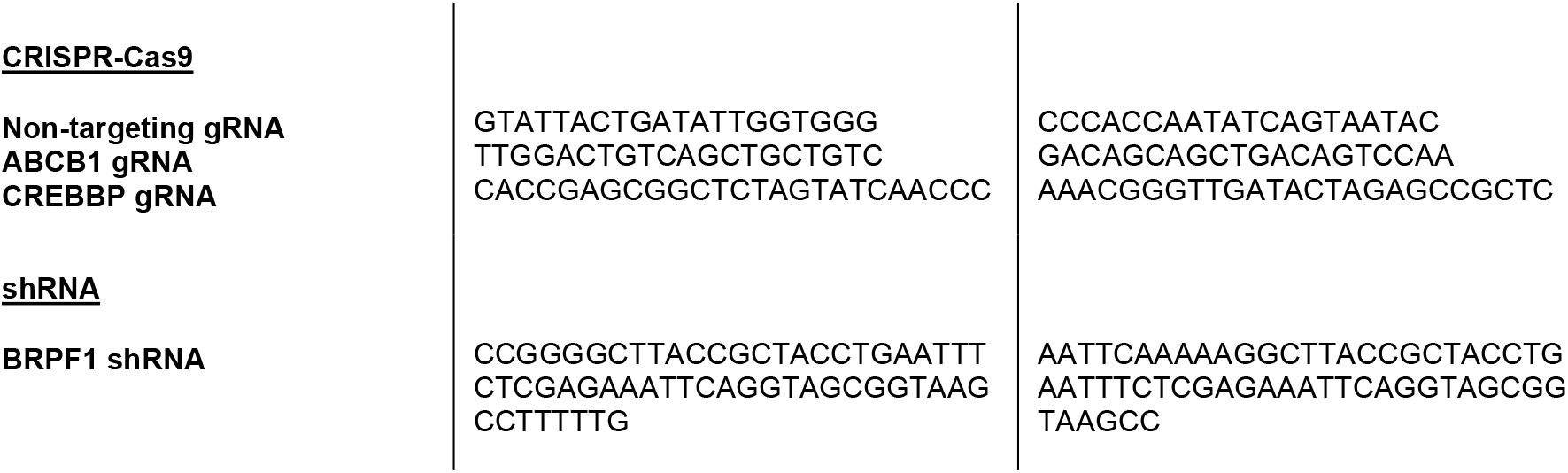
The nucleotide sequences of the primers used in the study.

**Sup. Table 2.** Top 10 ranked up- (red) and down-regulated (purple) genes in Du145-DtxR cells.

| Gene Name | LogFC | p-val | FDR | Description |
| --- | --- | --- | --- | --- |
| CP | 10.7 | 9.5E-107 | 2.5E-104 | Ceruloplasmin |
| ABCB1 | 9.1 | 3.7E-91 | 7.9E-89 | ATP Binding Cassette Subfamily B Member 1 |
| ABCC9 | 8.9 | 1.7E-12 | 1.7E-11 | ATP Binding Cassette Subfamily C Member 9 |
| KRTAP2-3 | 8.3 | 3.4E-96 | 7.6E-94 | Keratin Associated Protein 2-3 |
| C4orf19 | 7.6 | 4.6E-10 | 3.7E-09 | Chromosome 4 Open Reading Frame 19 |
| CFAP299 | 6.3 | 6.4E-07 | 3.5E-06 | Cilia And Flagella Associated Protein 299 |
| TDRD1 | 6.0 | 1.7E-06 | 9.2E-06 | Tudor Domain Containing 1 |
| GJA1 | 5.9 | 7.7E-101 | 1.8E-98 | Gap Junction Protein Alpha 1 |
| MIR4500HG | 5.7 | 1.7E-06 | 9.1E-06 | MIR4500 Host Gene |
| NFE4 | 5.7 | 1.9E-14 | 2.3E-13 | Nuclear Factor, Erythroid 4 |
| NUDT7 | -8.3 | 2.1E-11 | 1.9E-10 | Nudix hydrolase 7 |
| CA12 | -8.4 | 1.6E-11 | 1.5E-10 | Carbonic anhydrase 12 |
| TSNARE1 | -8.6 | 7.6E-12 | 7.3E-11 | t-SNARE domain containing 1 |
| SLC6A11 | -8.9 | 1.0E-12 | 1.0E-11 | Solute carrier family 6 member 11 |
| ADAMTS15 | -9.0 | 6.0E-13 | 6.3E-12 | ADAM metalloproteinase with thrombospondin type 1 motif 15 |
| COL5A1 | -9.0 | 4.8E-13 | 5.1E-12 | Collagen type V alpha 1 chain |
| S1PR3 | -9.5 | 2.6E-14 | 3.0E-13 | Sphingosine-1-phosphate receptor 3 |
| DAPK1 | -10.3 | 2.4E-16 | 3.3E-15 | Death associated protein kinase 1 |
| SH3BGRL | -11.2 | 6.0E-19 | 9.8E-18 | SH3 domain binding glutamate rich protein like |
| BCHE | -12.3 | 4.3E-22 | 8.8E-21 | Butyrylcholinesterase |

**Sup. Table 3.** Top 10 ranked up- (red) and down-regulated (purple) genes in Du145-CbzR cells.

| Gene Name | LogFC | p-val | FDR | Description |
| --- | --- | --- | --- | --- |
| ABCB1 | 12.0 | 1.1E-154 | 1.7E-151 | ATP Binding Cassette Subfamily B Member 1 |
| SLFN14 | 6.9 | 1.E-23 | 8.5E-22 | Schlafen Family Member 14 |
| HNRNPA1P9 | 6.7 | 2.4E-07 | 2.2E-06 | Heterogeneous Nuclear Ribonucleoprotein A1 Pseudogene 9 |
| CCDC187 | 6.0 | 2.0E-25 | 1.0E-23 | Coiled-Coil Domain Containing 187 |
| ABCB4 | 5.9 | 4.4E-15 | 1.0E-13 | ATP Binding Cassette Subfamily B Member 4 |
| MMP1 | 5.8 | 2.8E-89 | 1.2E-86 | Matrix Metalloproteinase 1 |
| ENSG00000286432 | 5.7 | 1.8E-23 | 8.1E-22 | lncRNA, novel transcript |
| MIR4500HG | 5.7 | 3.3E-06 | 2.5E-05 | MIR4500 Host Gene |
| MC5R | 5.4 | 8.8E-06 | 6.3E-05 | Melanocortin 5 Receptor |
| LINC01322 | 5.3 | 1.7E-43 | 2.2E-41 | Long Intergenic Non-Protein Coding RNA 1322 |
| EVC2 | -8.3 | 2.1E-11 | 1.9E-10 | EvC Ciliary Complex Subunit 2 |
| GPR173 | -8.3 | 1.6E-11 | 1.5E-10 | G Protein-Coupled Receptor 173 |
| DNAJC15 | -8.5 | 7.6E-12 | 7.3E-11 | DnaJ Heat Shock Protein Family (Hsp40) Member C15 |
| ZNF677 | -8.6 | 1.0E-12 | 1.0E-11 | Zinc Finger Protein 677 |
| TXNIP | -8.6 | 6.0E-13 | 6.3E-12 | Thioredoxin Interacting Protein |
| BRDT | -8.6 | 4.8E-13 | 5.1E-12 | Bromodomain Testis Associated |
| TINAGL1 | -10.3 | 2.6E-14 | 3.0E-13 | Tubulointerstitial Nephritis Antigen Like 1 |
| SLFN11 | -10.4 | 2.4E-16 | 3.3E-15 | Schlafen Family Member 11 |
| ZNF518B | -10.5 | 6.0E-19 | 9.8E-18 | Zinc Finger Protein 518B |
| BCHE | -12.8 | 4.3E-22 | 8.8E-21 | Butyrylcholinesterase |

**Sup. Table 4.**
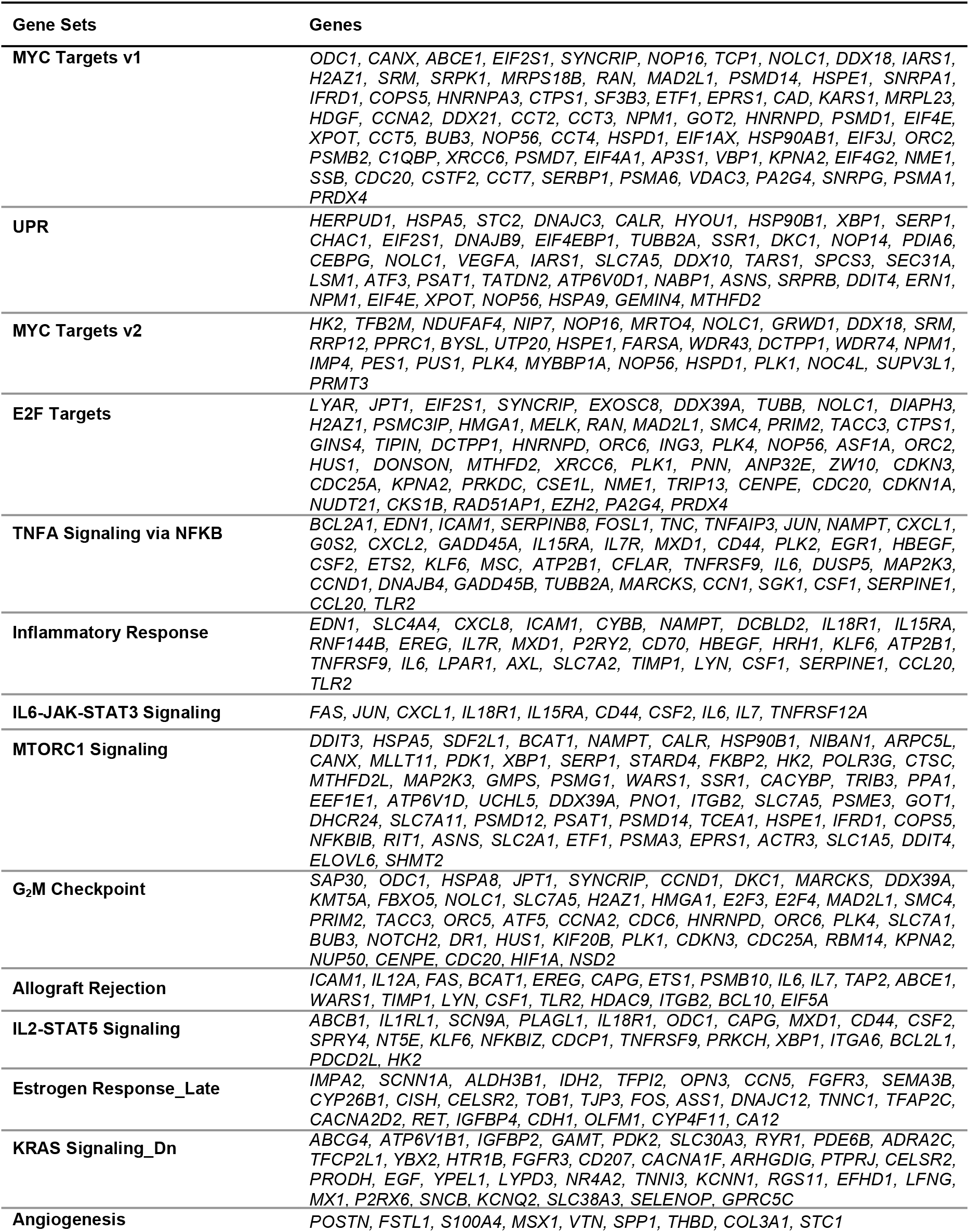
The genes in positively (NES ≥ 1.5) and negatively (NES ≤ 1.5) enriched gene sets in Du145-DtxR cells.

**Sup. Table 5.**
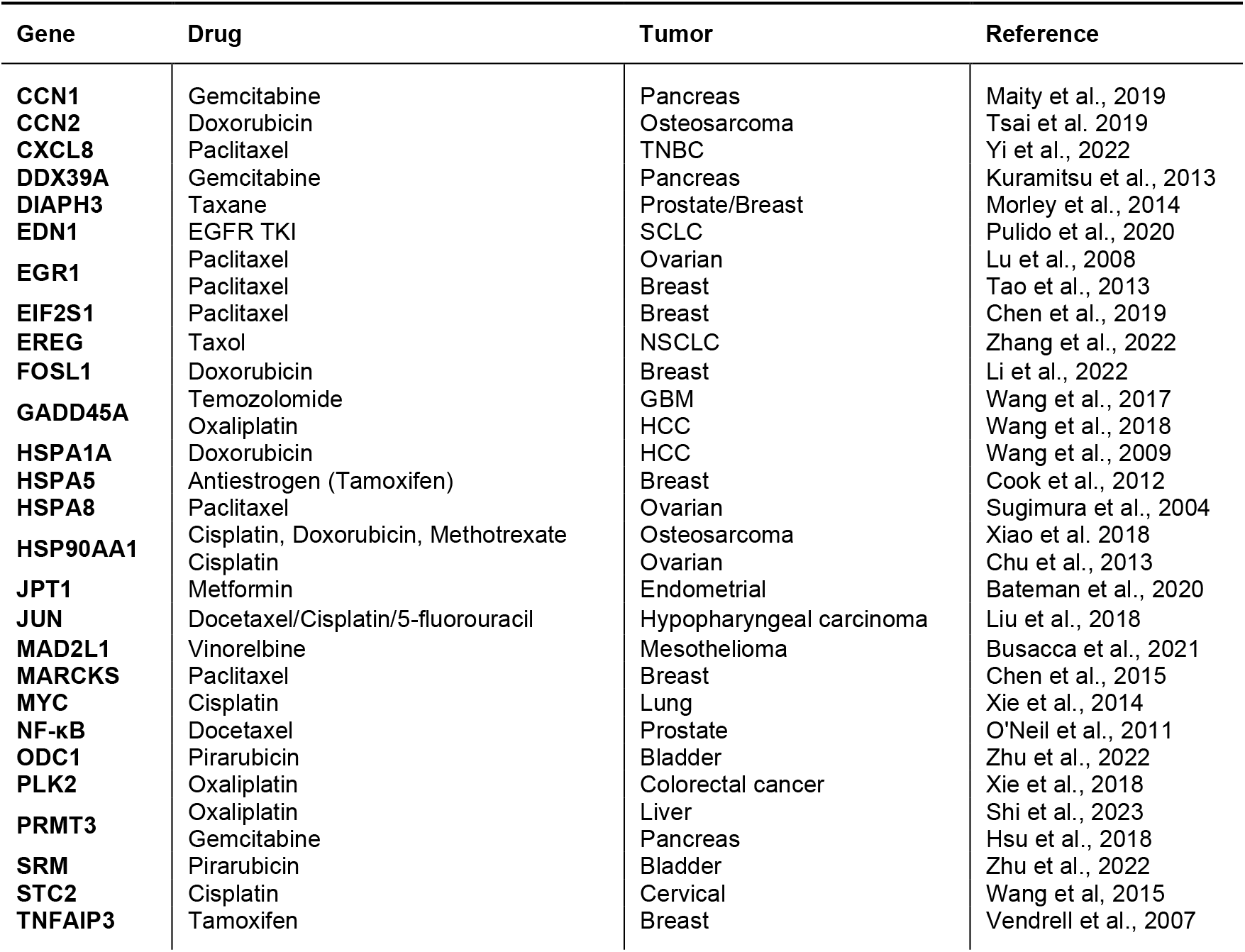
Relevance of the validated genes to drug resistance based on literature.

**Sup. Table 6.**
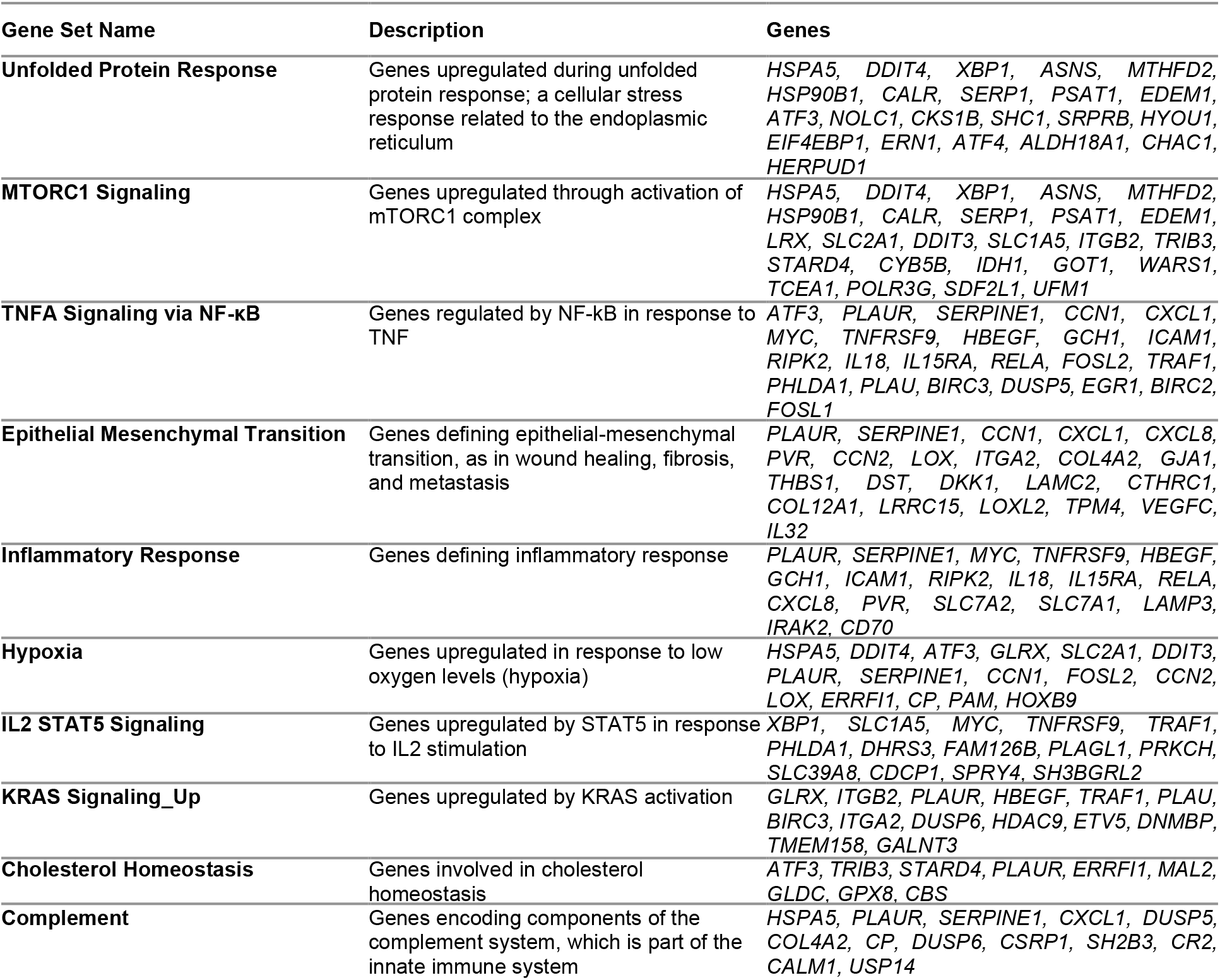
Representation of overlapping pathways/signaling of genes downregulated by siBRPF1 (The set of genes which lie in the intersection of the Venn diagram shown in **Figure 5G**.

**Sup. Table 7.** Representation of overlapping pathways/signaling of genes upregulated by siBRPF1 (The set of genes which lie in the intersection of the Venn diagram shown in **Sup. Figure 20A**.

| Gene Set Name | Description | Genes |
| --- | --- | --- |
| **Mitotic Spindle** | Genes important for mitotic spindle assembly | *FSCN1, TAOK2, ESPL1, RFC1, KIF15, BCL2L11, BIN1, DYNLL2, KLC1, TUBD1* |
| **Apical Junction** | Genes encoding components of apical junction complex | *FSCN1, TAOK2, PIK3R3, PTEN, MAP3K20, B4GALT1, GNAI2, CTNND1, LDLRAP1, PBX2* |
| **E2F Targets** | Genes encoding cell cycle related targets of E2F transcription factors | *ESPL1, RFC1, PPM1D, STMN1, CDC25B, MCM2, RAD21, NUP205, MXD3, TIMELESS* |
| **PI3K-AKT-mTOR Signaling** | Genes up-regulated by activation of the PI3K/AKT/mTOR pathway | *PIK3R3, PTEN, MYD88, AKT1, RAC1, UBE2N, DUSP3* |
| **IFNγ Response** | Genes up-regulated in response to IFNG [GeneID=3458]. | *MYD88, IFIT2, OGFR, IFI35, IFIT3, IFIT1, TNFAIP2, LATS2, SPPL2A* |
| **P53 Pathway** | Genes involved in p53 pathways and networks | *PPM1D, S100A10, RXRA, ACVR1B, S100A4, DDB2, MXD4, HEXIM1, PRMT2* |
| **G<sub>2</sub>M Checkpoint** | Genes involved in the G<sub>2</sub>/M checkpoint, as in progression through the cell division cycle | *ESPL1, KIF15, MAP3K20, STMN1, CDC25B, MCM2, RAD21, E2F2* |
| **Estrogen Response_Late** | Genes defining late response to estrogen. | *CCN5, BAG1, NBL1, CYP26B1, ACOX2, MEST, IMPA2* |
| **IFNα Response** | Genes up-regulated in response to alpha interferon proteins | *IFIT2, OGFR, IFI35, IFIT3, CNP* |
| **Androgen Response** | Genes defining response to androgens | *B4GALT1, AKT1, KRT8, H1-0, APPBP2* |

**Sup. Figure 1.**
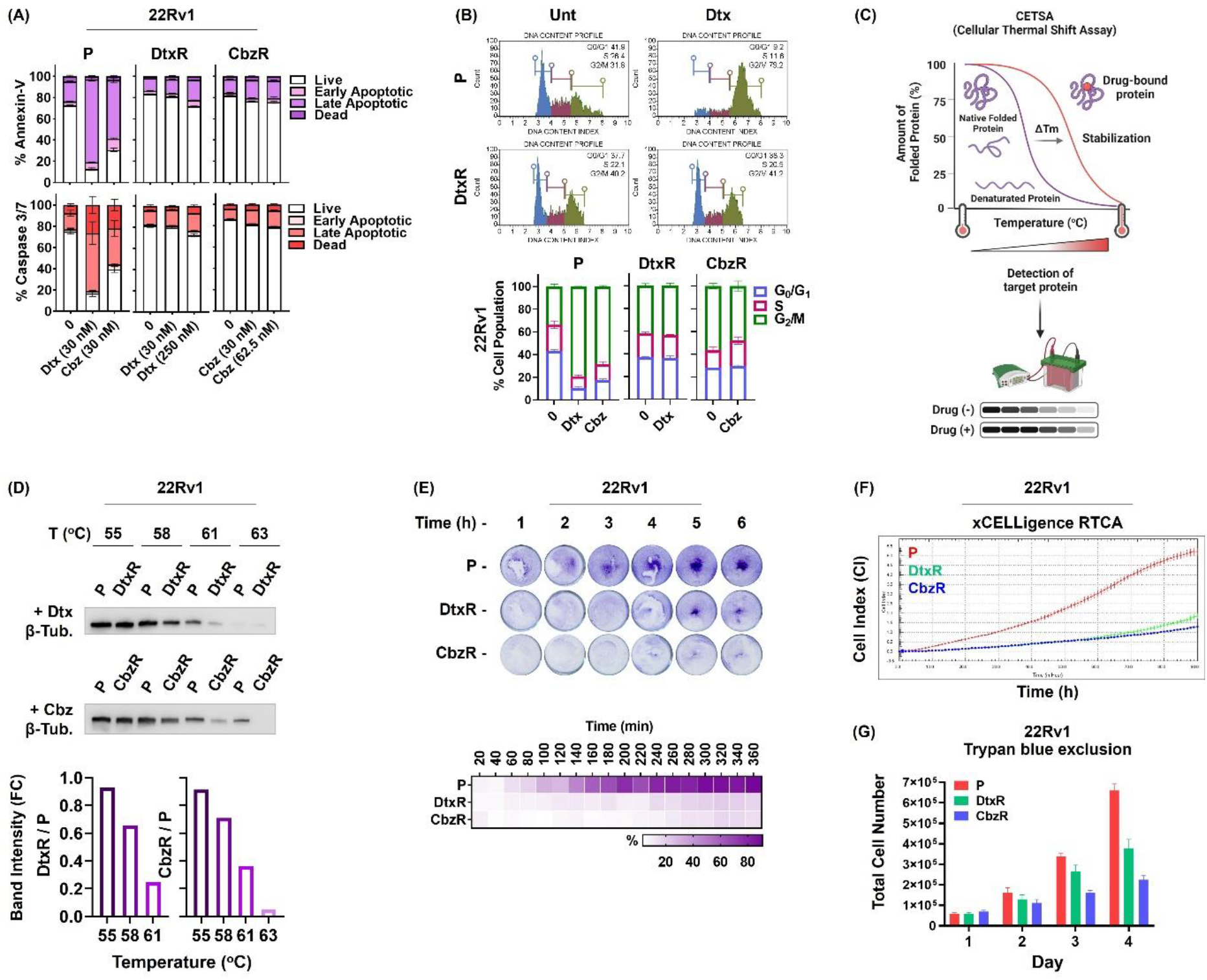
**(A)** Flow cytometry analysis of cell death in taxane treated 22Rv1-P and -R cells. **(B)** The cell cycle distribution of 22Rv1-P/R cells treated with or without Dtx/Cbz (15 nM). Representative DNA content histograms (upper panel) and quantifications are shown (lower panel). **(C)** Diagram showing the principle of CETSA. Figure was made in BioRender. **(D)** CETSA for in-cell β-Tubulin target engagement. Western blots showing thermostable β-Tubulin following indicated heat shocks in the presence of Dtx (62.5 nM) or Cbz (31.5 nM) in Du145-P/R cells. CETSA images were quantified by using ImageJ software (shown below). **(E)** Adhesion capacity of 22Rv1-P/R cells. Attached cells were fixed at the indicated time points and were stained with crystal violet. Quantification was performed using ImageJ software (shown below). **(F)** Real time monitoring of cell growth of 22Rv1-P/R cells with the xCELLigence system. **(G)** The growth rate of 22Rv1-P/R was also determined by counting live cells by trypan blue staining. **P:** parental, **R:** resistant.

**Sup. Figure 2.**
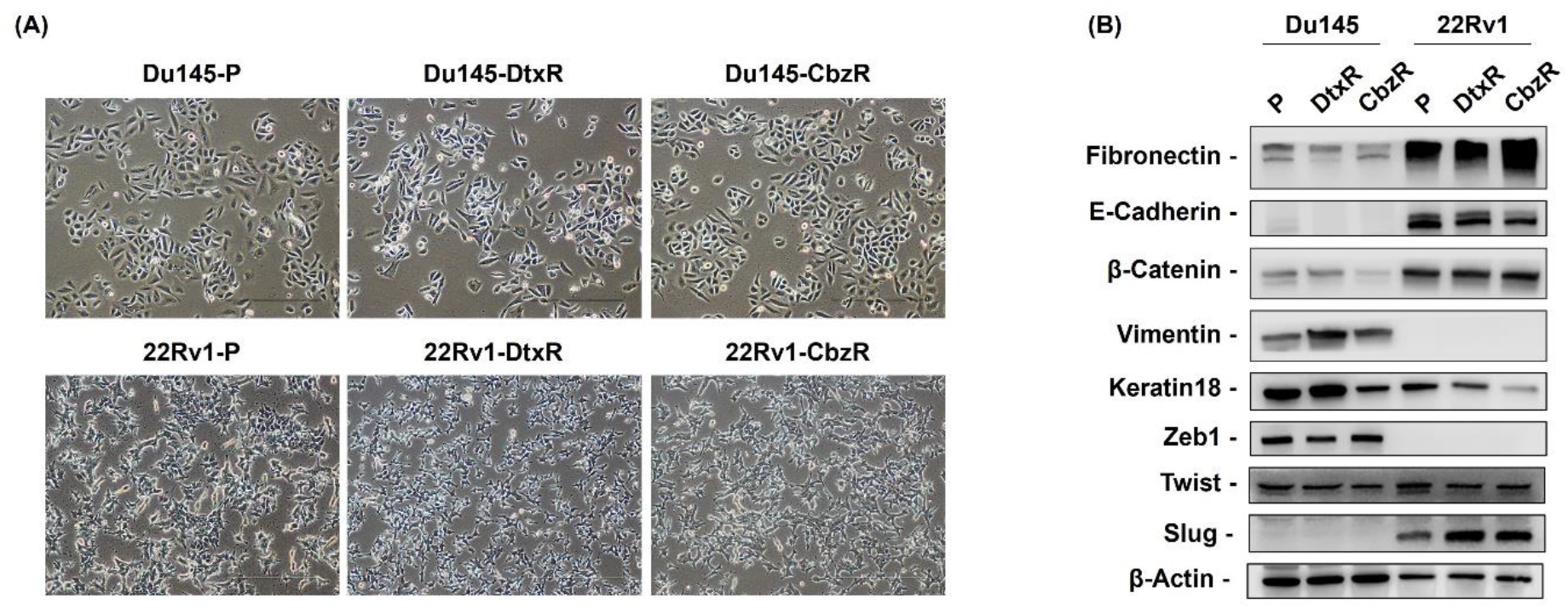
**(A)** Cell morphology of parental and resistant PCa (Du145 and 22Rv1) cells. Images were acquired by using a phase contrast microscope (magnification, ×100). **Dtx:** Docetaxel. **Cbz:** Cabazitaxel. **(B)** Western blot analysis of EMT markers in parental and resistant PCa cells. **P:** parental, **R:** resistant.

**Sup. Figure 3.**
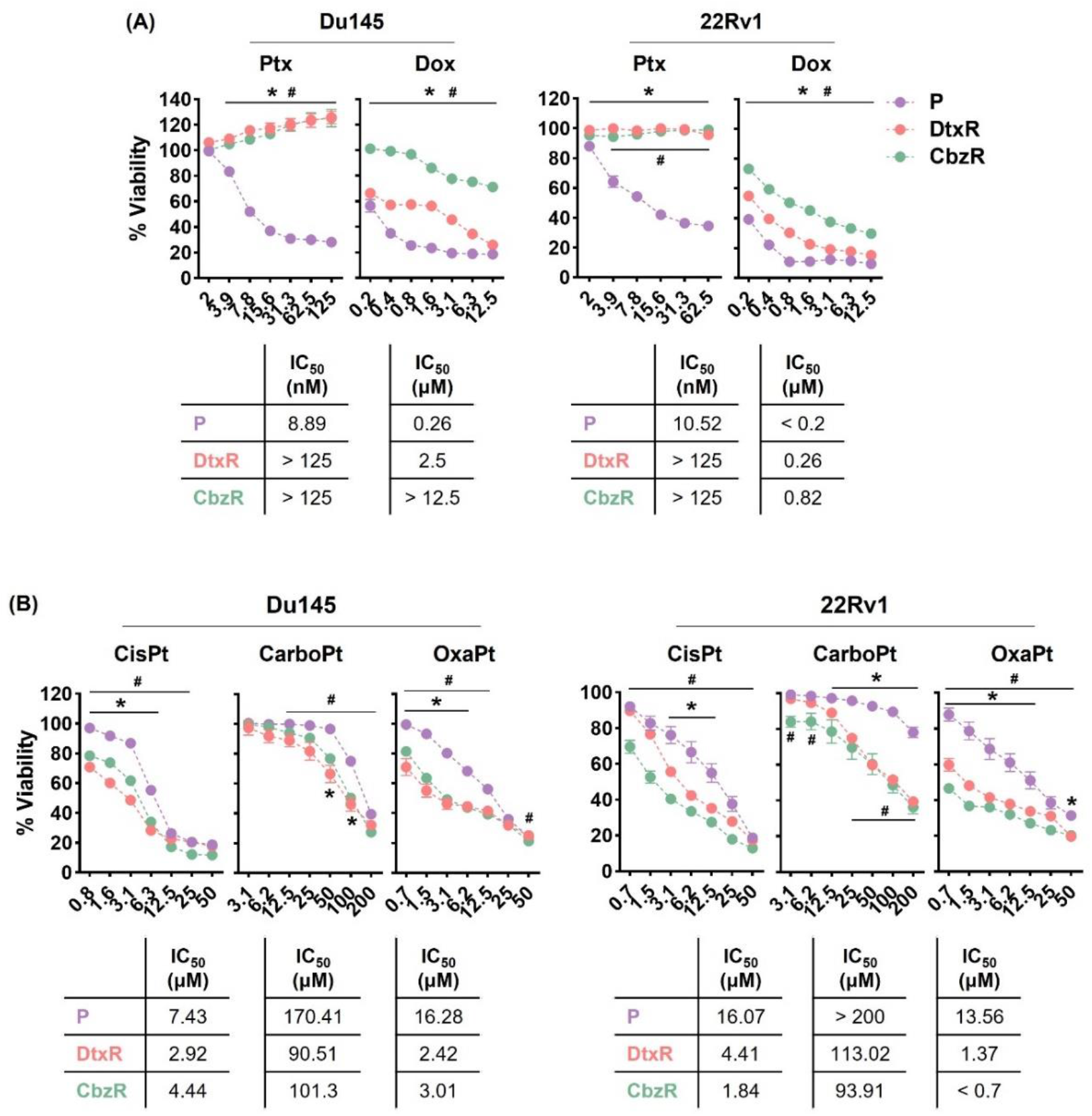
The dose dependent effects of **(A)** paclitaxel (Ptx), doxorubicin (Dox), **(B)** cisplatin (CisPt), carboplatin (CarboPt) and oxaliplatin (OxaPt) on the cell viability of parental and resistant PCa cells. The results were obtained by using SRB assay (72 h) and expressed as the mean ± SEM of triplicate from 2 biological replicates. The IC_50_ value of each cell line is shown in corresponding colors below the dose-response curves. (*) and (#) indicate significant differences in cell viability between parental and DtxR or CbzR cells, respectively (p < 0.05). **P:** parental, **R:** resistant.

**Sup. Figure 4.**
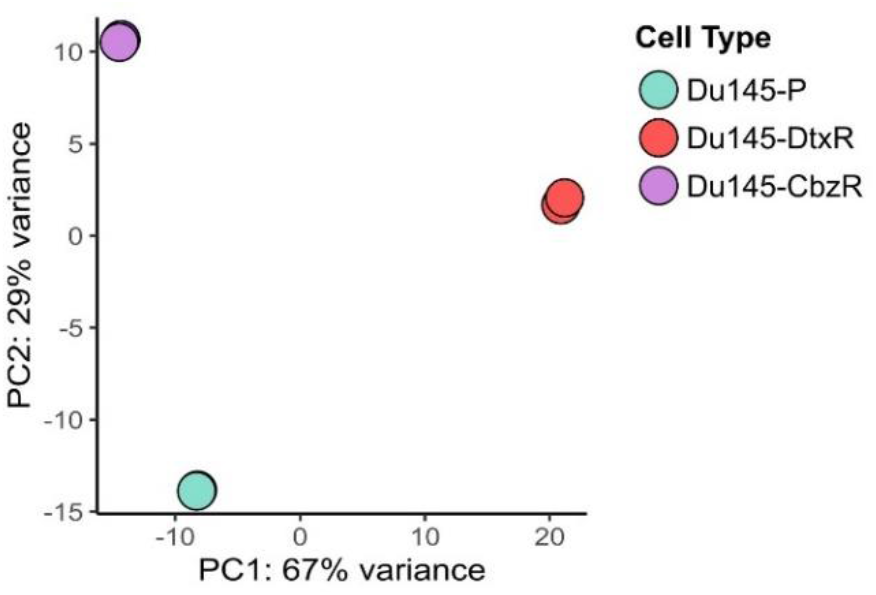
PCA plot of RNA-seq samples from parental (P), Docetaxel- (DtxR) and Cabazitaxel-resistant (CbzR) Du145 cells.

**Sup. Figure 5.**
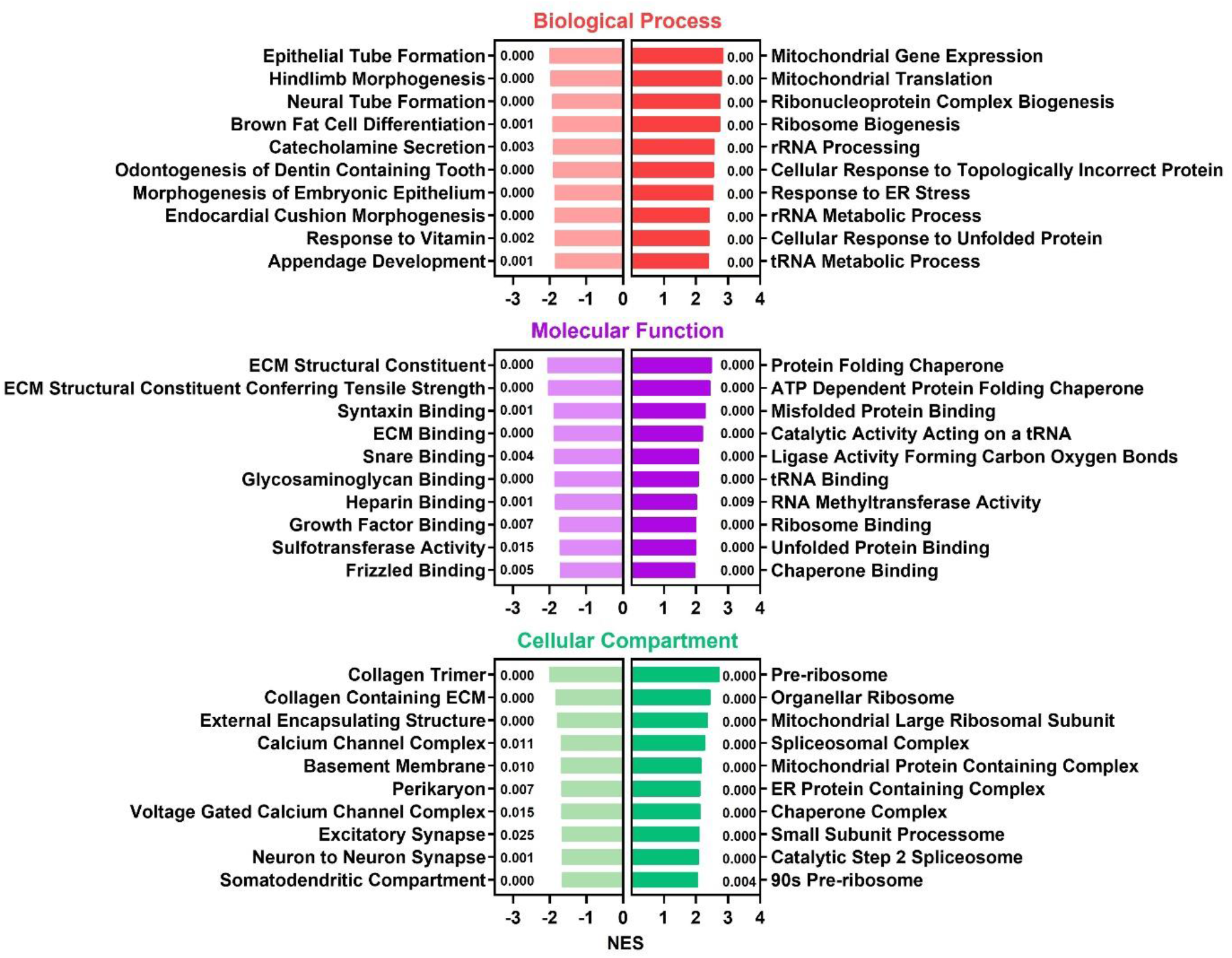
Top 10 Gene ontology (GO) enrichment analysis of DEGs between Du145-DtxR and Du145-P cells (FDR < 0.05, and Log_2_FC ≥ 0.5 or ≤ -0.5). The numbers next to the columns indicate the NOM p-val values. GSEA was performed using c5.go.bp.v2023.1.Hs.symbols.gmt, c5.go.mf.v2023.1.Hs.symbols.gmt, and c5.go.cc.v2023.1.Hs.symbols.gmt datasets in the MsigDB database.

**Sup. Figure 6.**
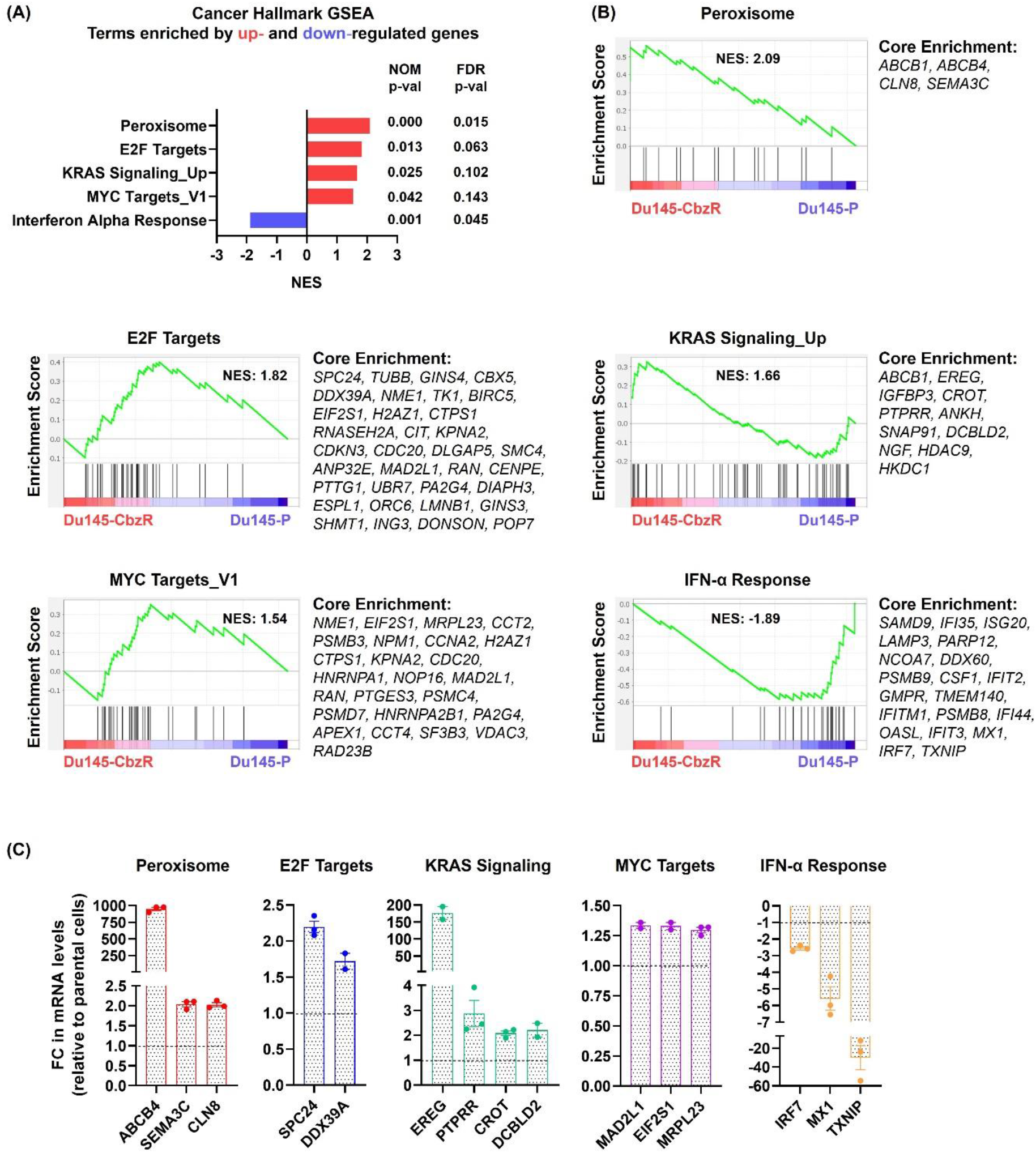
**(A)** Gene set enrichment (GSEA) analysis using hallmark gene sets from the MSigDB in Du145-DCbzR cells (FDR < 0.05, and Log_2_FC ≥ 0.5 or ≤ -0.5). **(B)** Enrichment plots for the gene sets enriched in GSEA Hallmark analysis. **(C)** Validation of enriched core genes. Expression levels of mRNAs were determined by qRt-PCR. Data is the mean ± SEM.

**Sup. Figure 7.**
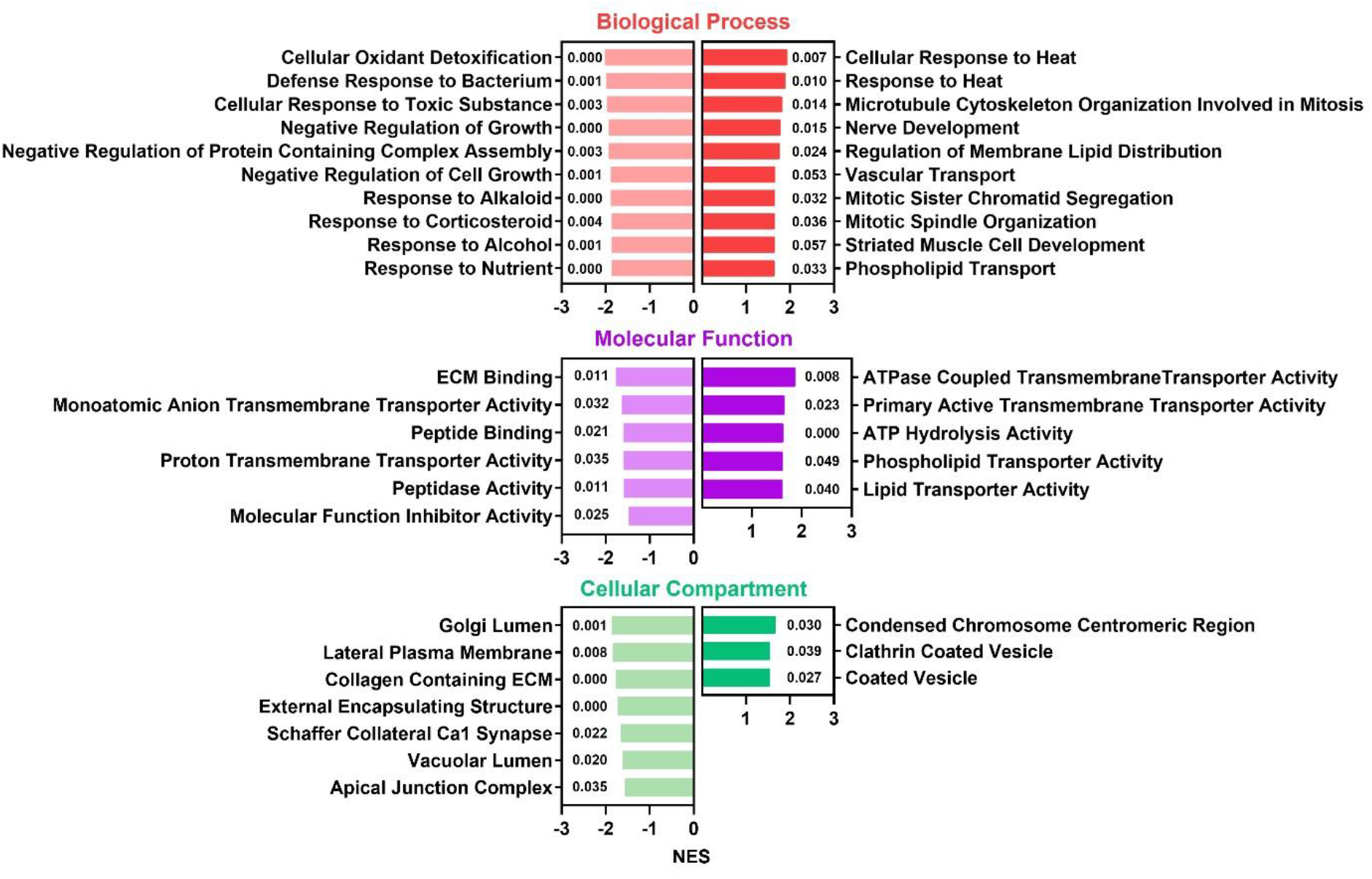
Top 10 Gene ontology (GO) enrichment analysis of DEGs between Du145-CbzR and Du145-P cells (FDR < 0.05, and Log_2_FC ≥ 0.5 or ≤ -0.5). The numbers next to the columns indicate the NOM p-val values. GSEA was performed using c5.go.bp.v2023.1.Hs.symbols.gmt, c5.go.mf.v2023.1.Hs.symbols.gmt, and c5.go.cc.v2023.1.Hs.symbols.gmt datasets in the MsigDB database.

**Sup. Figure 8.**
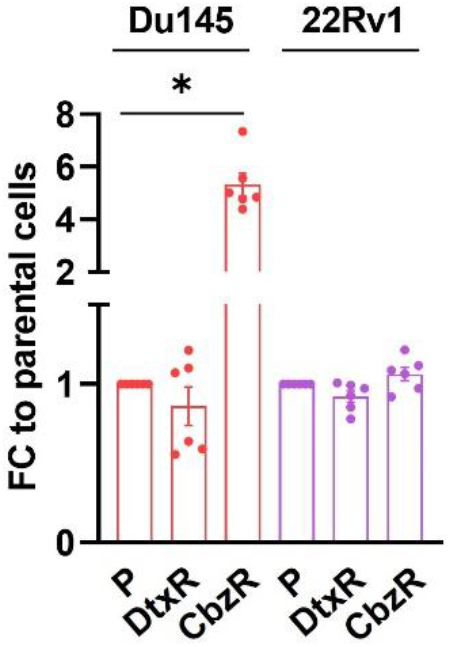
ABCB1 amplification in taxane resistant PCa cells. The data expressed as mean ± SEM from 2 biological replicates. (*) indicates significant differences ABCB1 gene copy numbers between parental and resistant cells (p < 0.0001).

**Sup. Figure 9.**
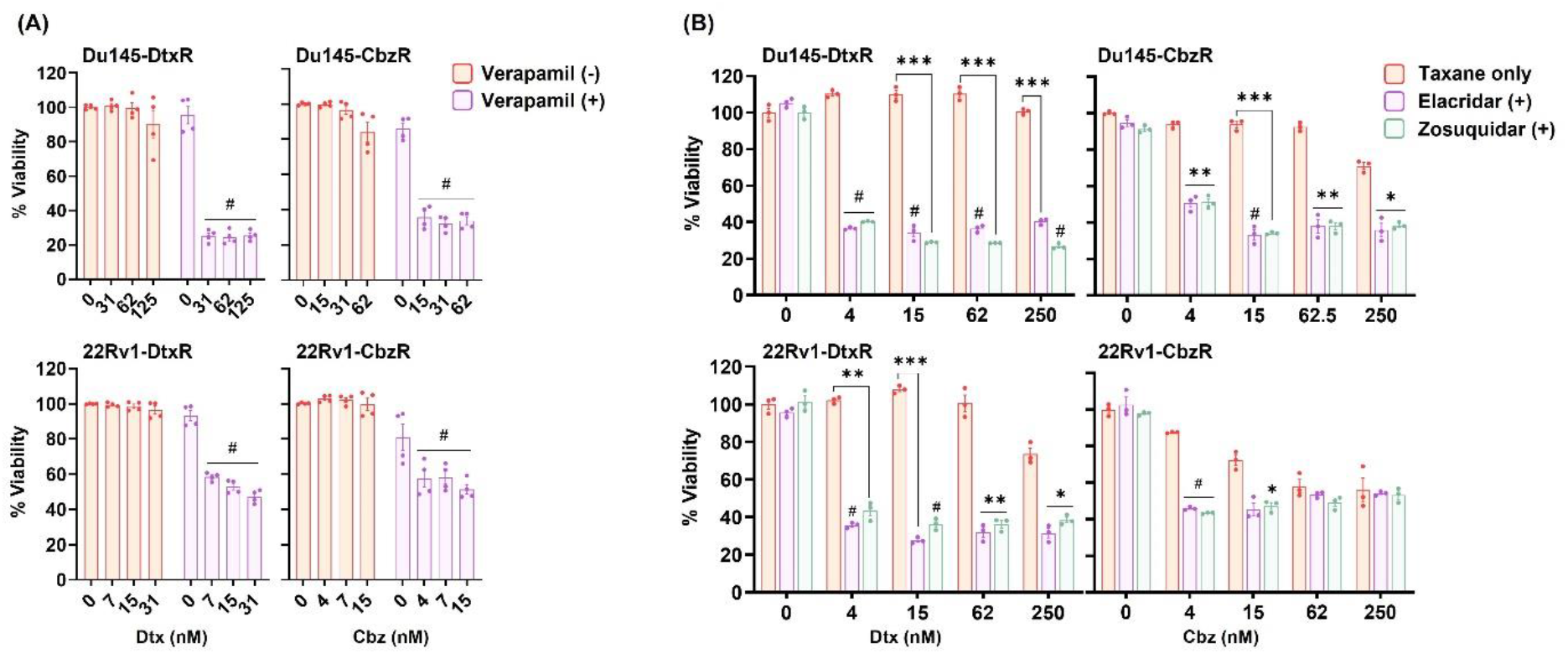
Dtx and Cbz resistance were reversible following ABCB1 interference. SRB viability assay was used to compare cell viabilities (72 h) following exposure to serial dilutions of Dtx and Cbz in the presence of ABCB1-inhibitors; **(A)** verapamil (20 µM), **(B)** elacridar and zosuquidar (0.5 - 0.12 µM). The symbols * (p < 0.05), ** (p < 0.01), *** (p < 0.001), and # (p < 0.0001) indicate the statistical significance in cell viability between ABCB1-inhibitor treated groups and non-treated groups.

**Sup. Figure 10.**
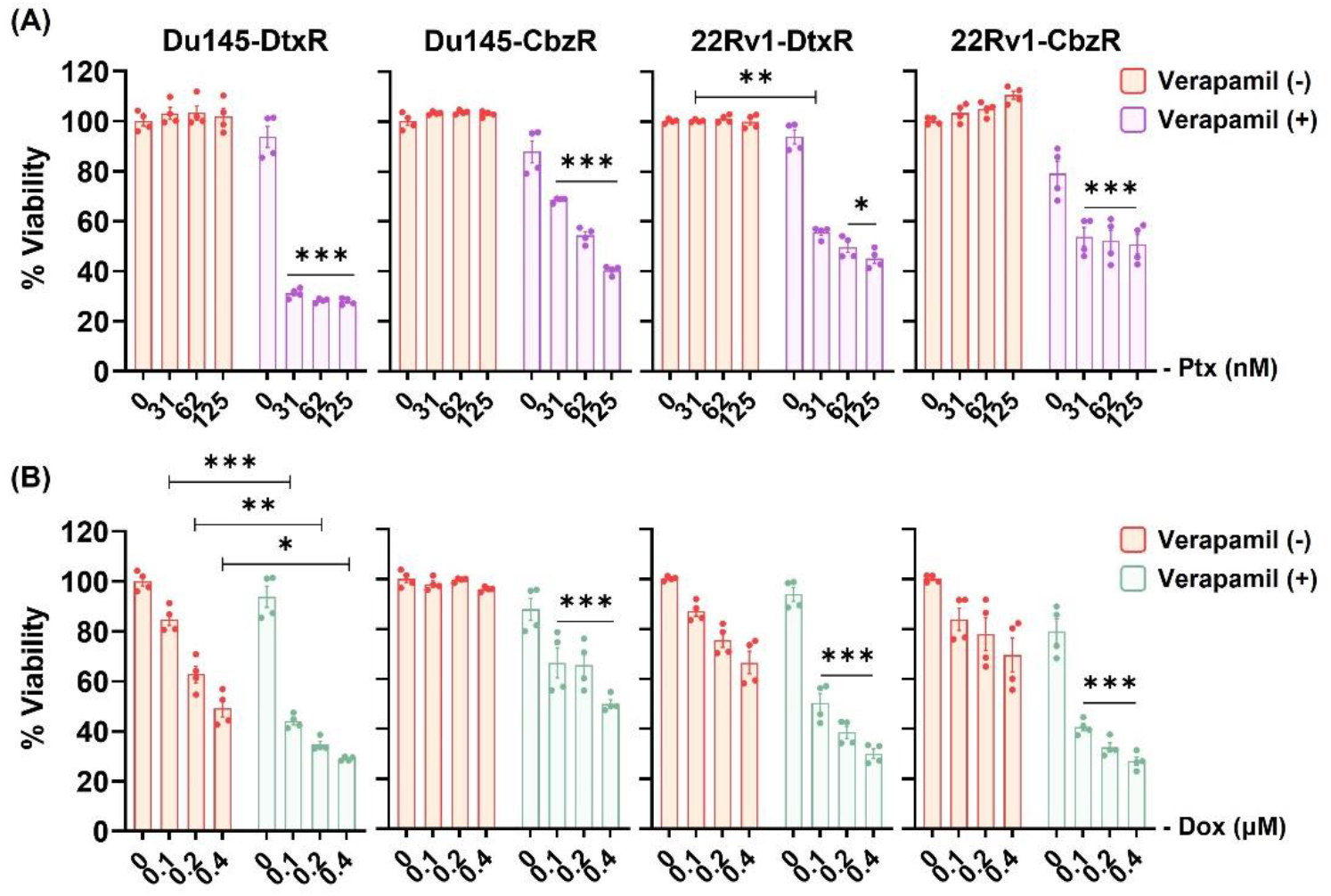
Cell viabilities of drug resistant cells following exposure to Paclitaxel (Ptx) and Doxorubicin (Dox) in the presence of verapamil (20 µM). The results were obtained by SRB assay (72h) and expressed as mean ± SEM from 2 biological replicates. The symbols (*) p < 0.01, (**) p < 0.001, and (***) p < 0.001 indicate significant differences in cell viability between verapamil treated groups and non-treated groups.

**Sup. Figure 11.**
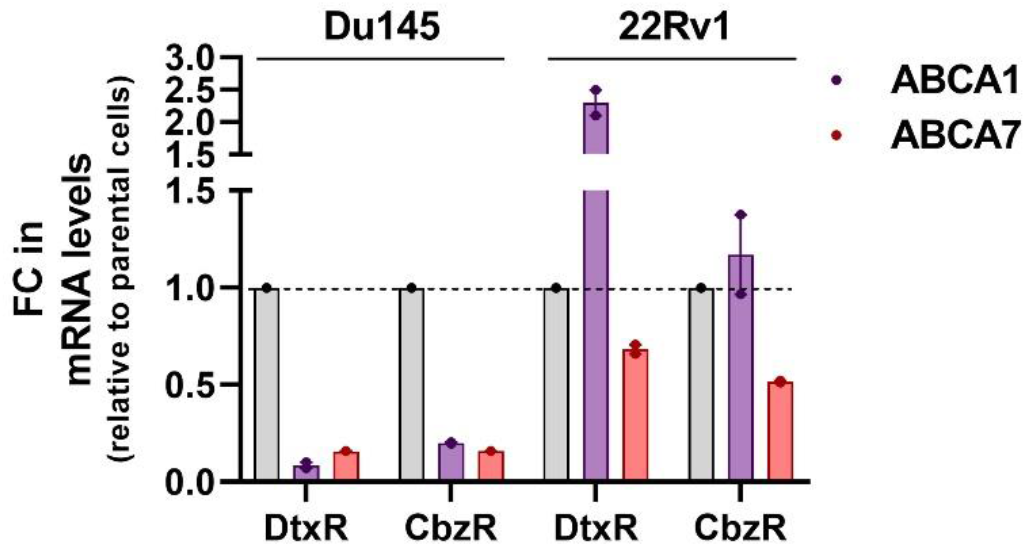
ABCA1 and ABCA7 mRNA expression levels were determined by using qRT-PCR. Data is the mean ± SEM. The fold change (FC) values were calculated relative to the parental cells (gray columns).

**Sup. Figure 12.**
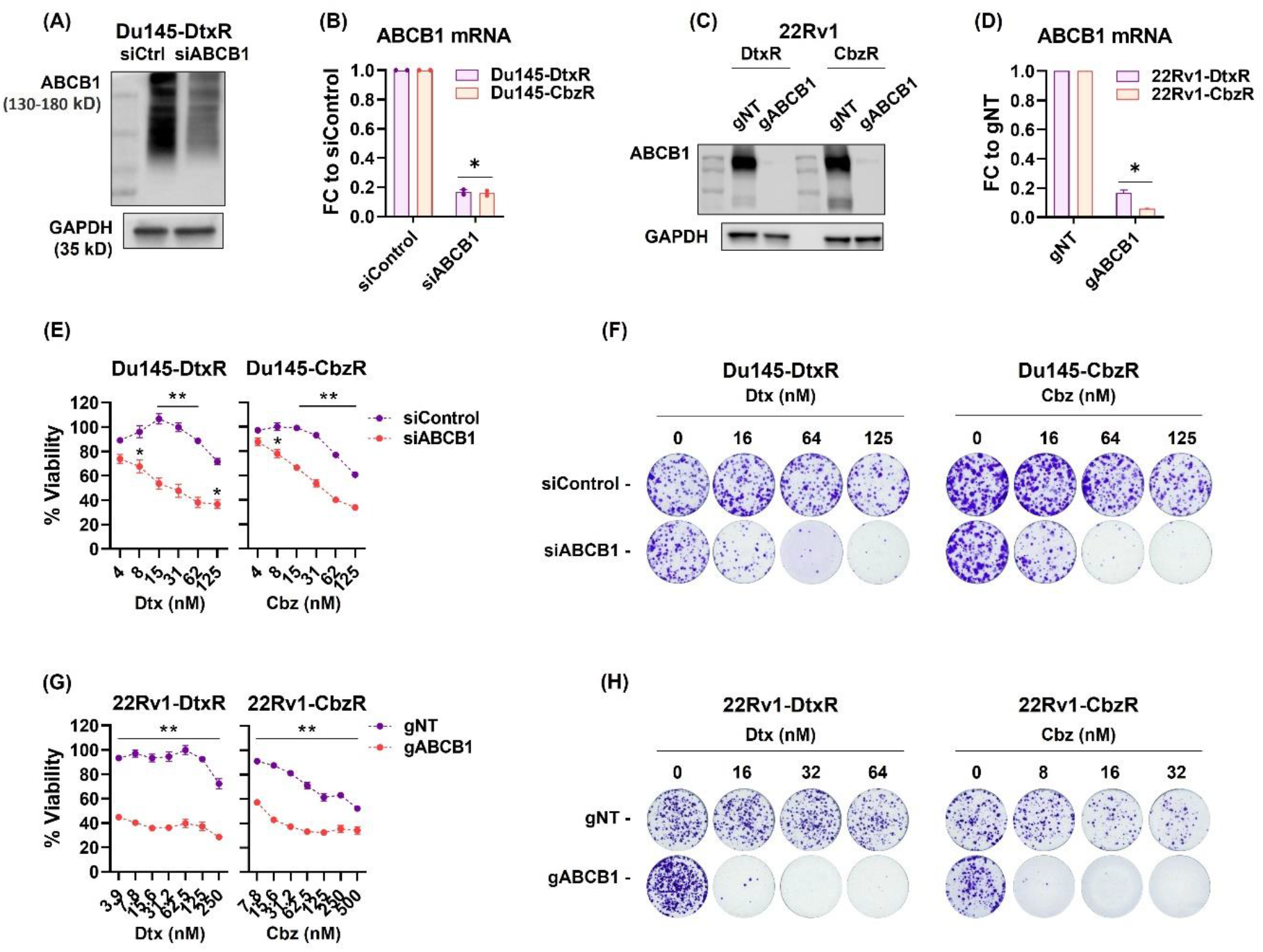
The efficiency of siRNA mediated ABCB1 knockdown was evaluated by performing both **(A)** western blot and **(B)** qRt-PCR analyses. **(C)** The efficiency of ABCB1 knockout (CRISPR-Cas9) in 22Rv1-R cells was validated using Western blot and **(D)** qRt-PCR analysis. **(E, F)** Taxane response in ABCB1 silenced cells was assessed by SRB (72h) and clonogenic viability assays. **(G, H)** SRB (72h) and clonogenic viability assays were performed on gABCB1 and gNT treated 22Rv1-R cells. The data expressed as mean ± SEM. Statistical significance denoted as (*) p < 0.001 and (**) p < 0.0001.

**Sup. Figure 13.**
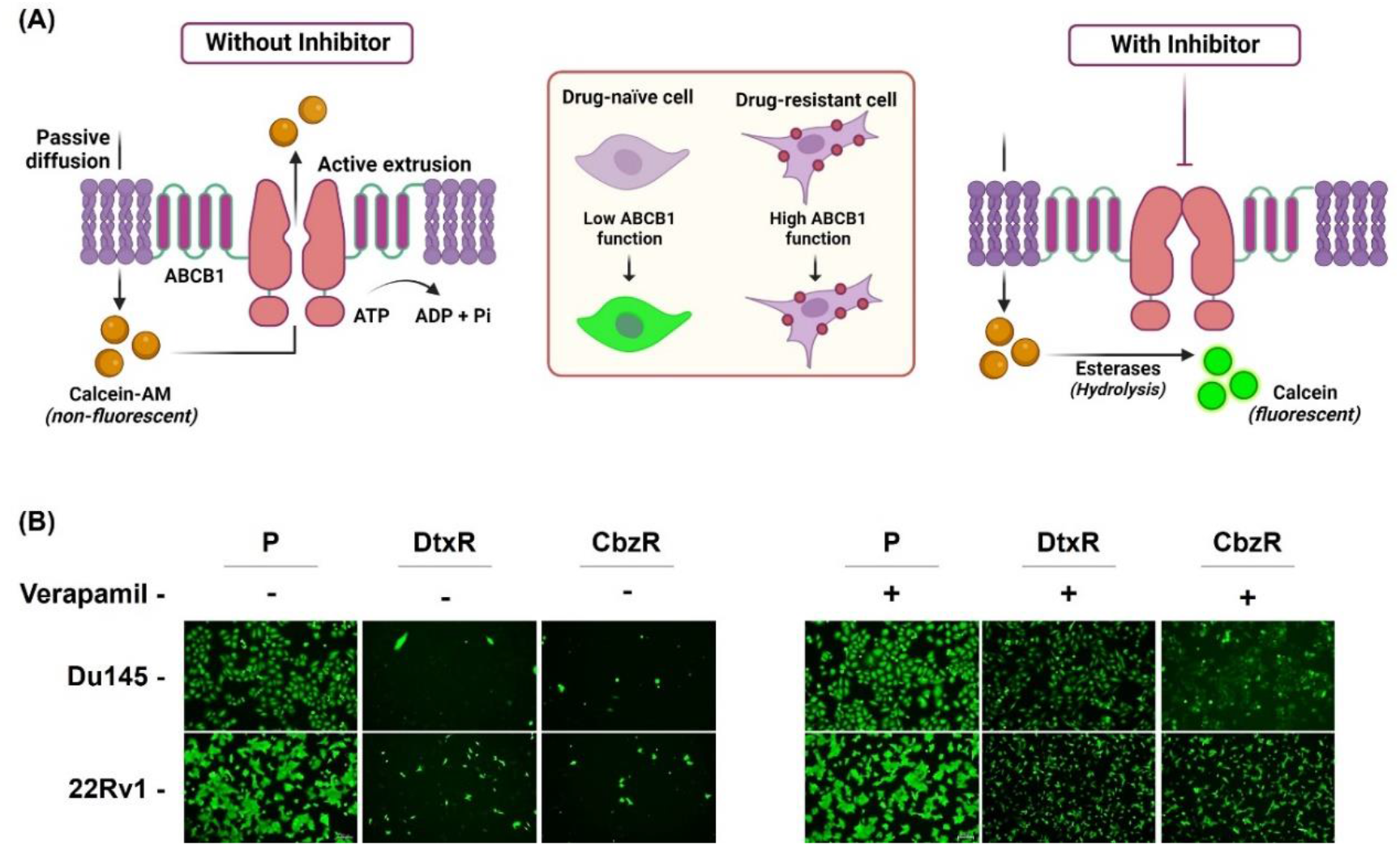
**(A)** Diagram showing the principle of the calcein efflux assay. Figure was made in BioRender. **(B)** Parental and resistant PCa cells were incubated with calcein-AM in the absence (left panel) or presence (right panel) of verapamil (20 µM, 8h) and visualized under a fluorescence microscope.

**Sup. Figure 14.**
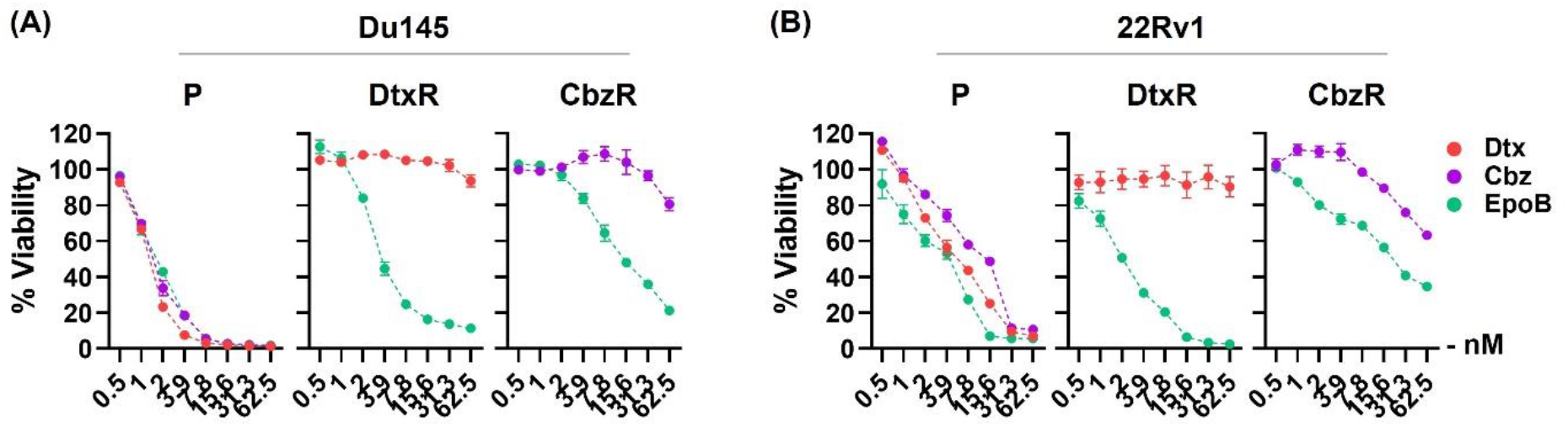
Dose dependent effects of Epothilone B (Epo B), Dtx or Cbz on the cell viability of parental and taxane resistant **(A)** Du145 and **(B)** 22Rv1 cells. The results were obtained by CTG assay (72h) and expressed as mean ± SEM. **P:** parental, **R:** resistant.

**Sup. Figure 15.**
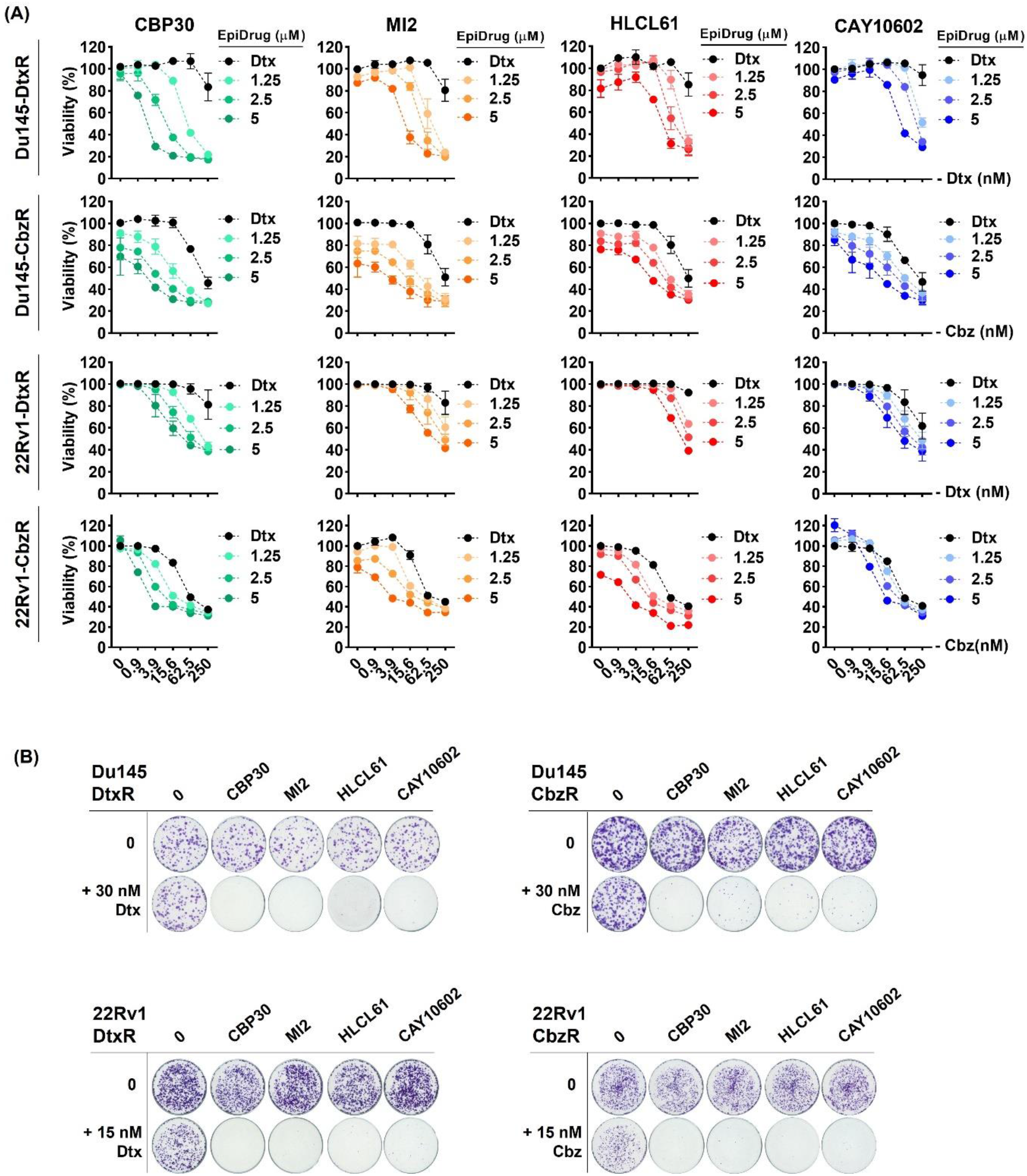
Validation dose-response of epidrugs (CBP30, MI2, HLCL61 and CAY10602) on taxane resistant PCa cells. **(A)** Cells were co-treated with indicated drugs (taxane; 1-250 nM and indicated inhibitors; 1.25-5 µM) and the results were obtained by SRB viability assay (72 h). The data is expressed as mean ± SEM. **(B)** Clonogenic images were obtained by treating cells with indicated drugs for 72 h and the colony formation ability was analyzed 10-15 days after drug exposure.

**Sup. Figure 16.**
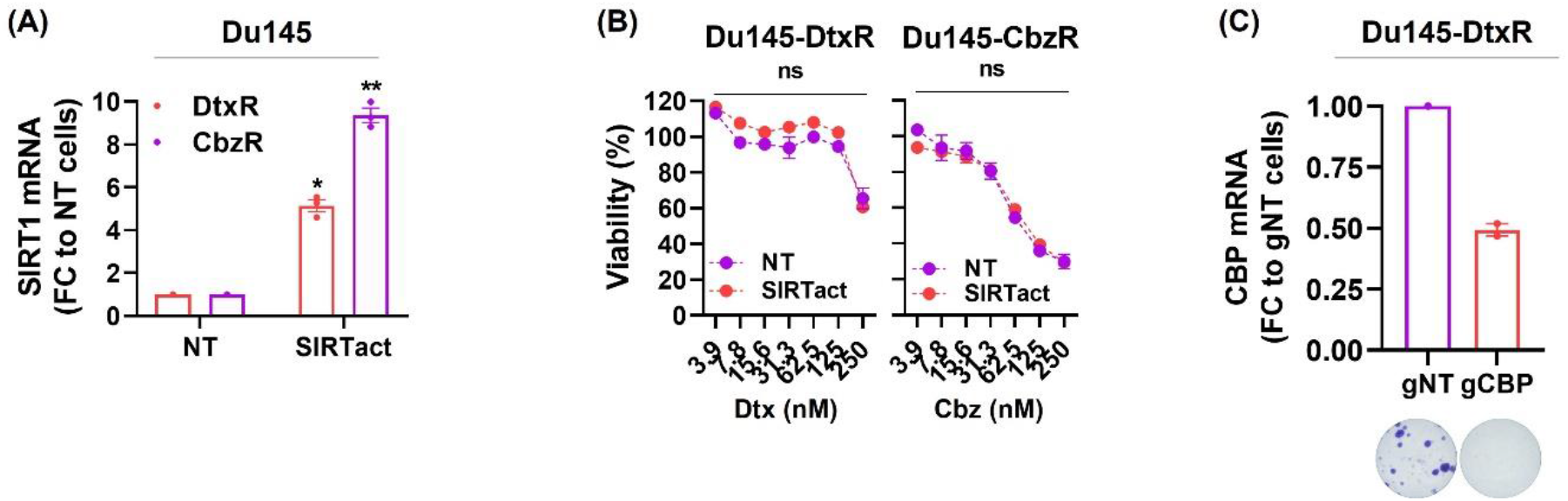
**(A)** Validation of SIRT overexpression by qRT-PCR, comparing non-transduced (NT) cells to stably SIRT1 expressing counterparts. The data is expressed as mean ± SEM. Statistical significance denoted as (*) p < 0.01 and (**) p < 0.001. **(B)** Taxane response of SIRT1-expressing cells was assessed by SRB viability assay (72h). **(C)** Knockdown efficiency of gCBP in Du145-DtxR cells. After transduction with control (gNT) and CBP-targeting gRNAs, the expression of CBP was detected by qRt-PCR. The colony formation ability of CBP-guided cells is represented under the corresponding column.

**Sup. Figure 17.**
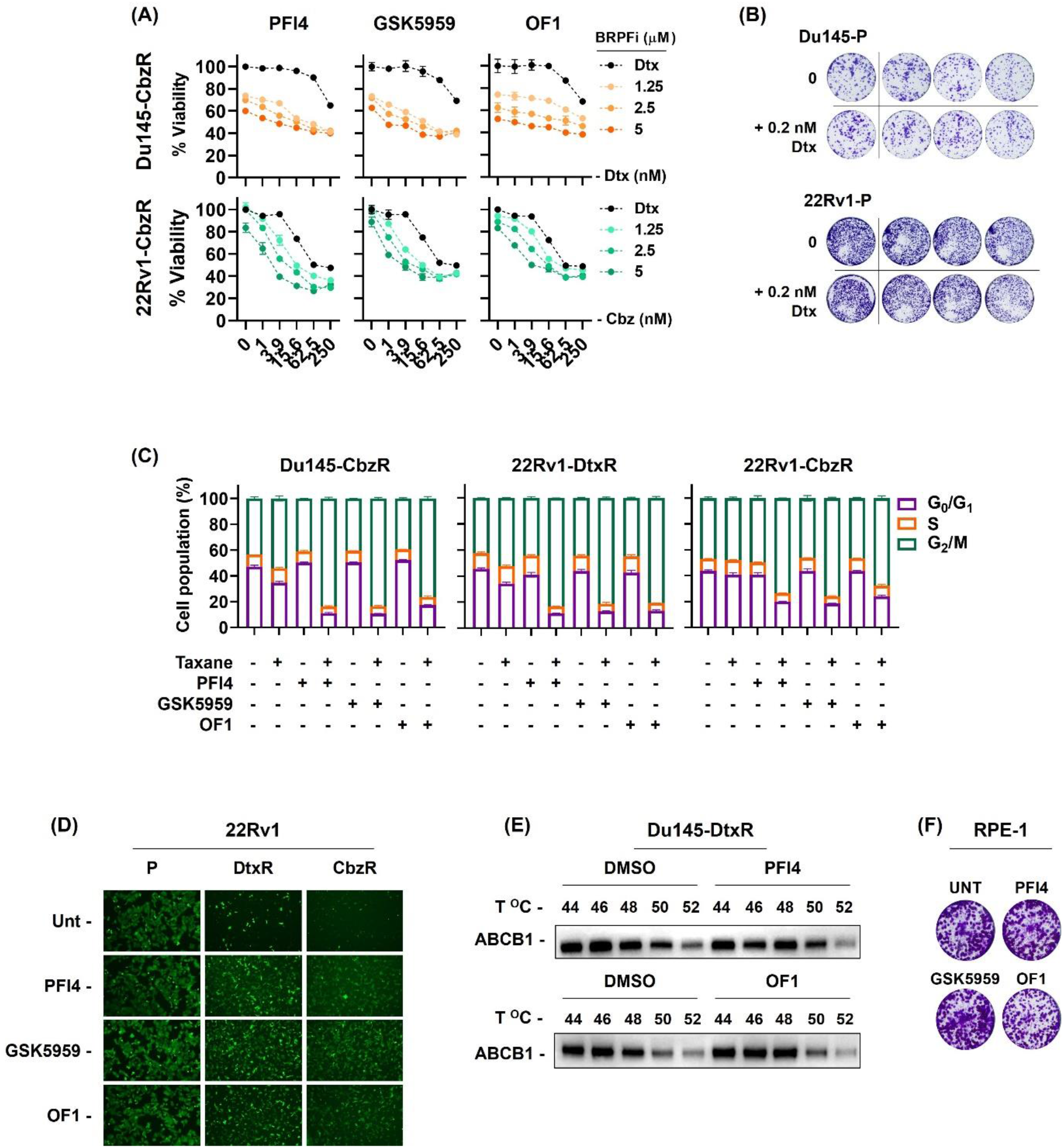
**(A)** Validation dose-response curves of BRPF inhibitors (PFI4, GSK5959 and OF1) on Cbz-resistant cells. Cells were co-treated with indicated drugs (Dtx; 1-250 nM and BRPF inhibitors; 1.25-5 µM) and the results were obtained by SRB viability assay (72 h). The data is expressed as mean ± SEM. **(C)** Cell cycle distribution (24h) in resensitized resistant cells. **(D)** Calcein retention assay was performed in the absence or presence of BRPF inhibitors (5 µM, 24h). **(E)** CETSA for in-cell ABCB1 engagement. Western blots showing thermostable ABCB1 following indicated heat shocks (44°C, 46°C, 48°C, 50°C and 52°C) in the presence of the indicated BRPF inhibitors (5 µM) in Du145-DtxR cells. **(F)** The efficacy of BRPF inhibitors on RPE-1 cells was evaluated using colony formation assay.

**Sup. Figure 18.**
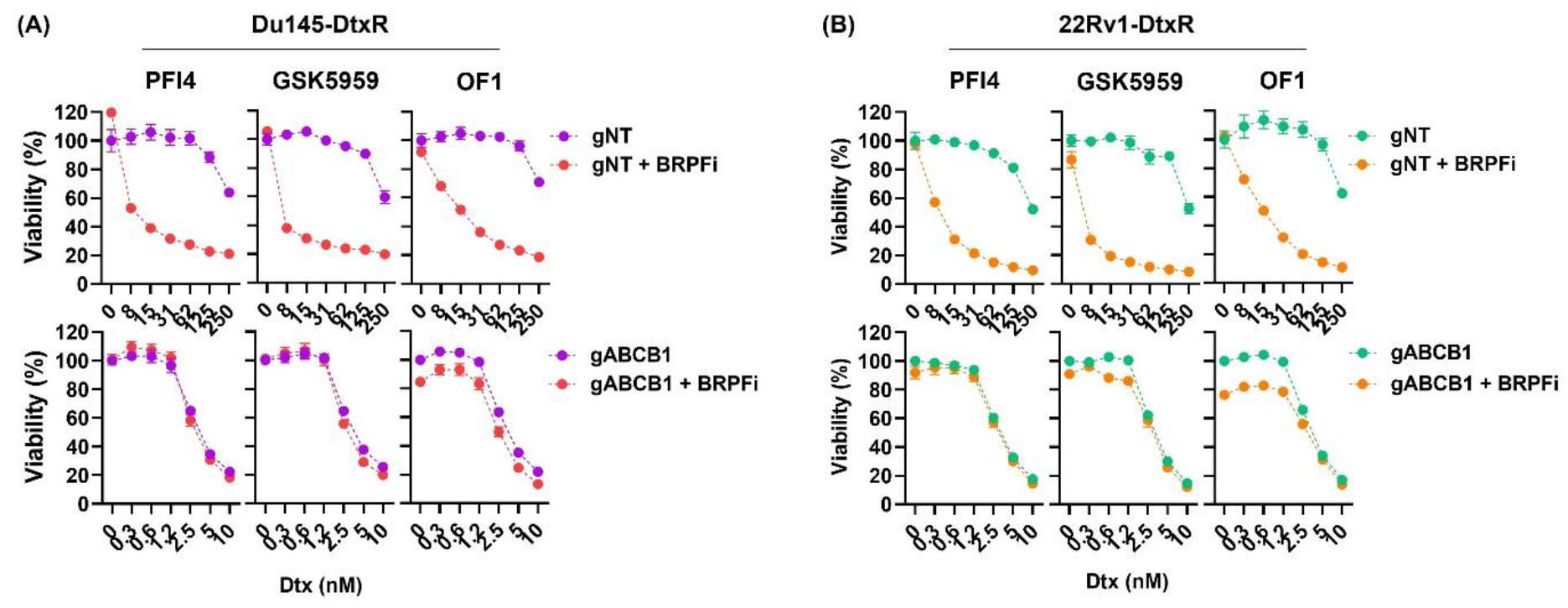
Viability graphs of non-targeting (NT) and ABCB1-targeting (gABCB1) guide received **(A)** Du145-DtxR and **(B)** 22Rv1-DtxR cells co-treated with BRPF inhibitors (5 µM) and Dtx. The results were obtained by CTG viability assay (72 h). The data is expressed as mean ± SEM.

**Sup. Figure 19.**
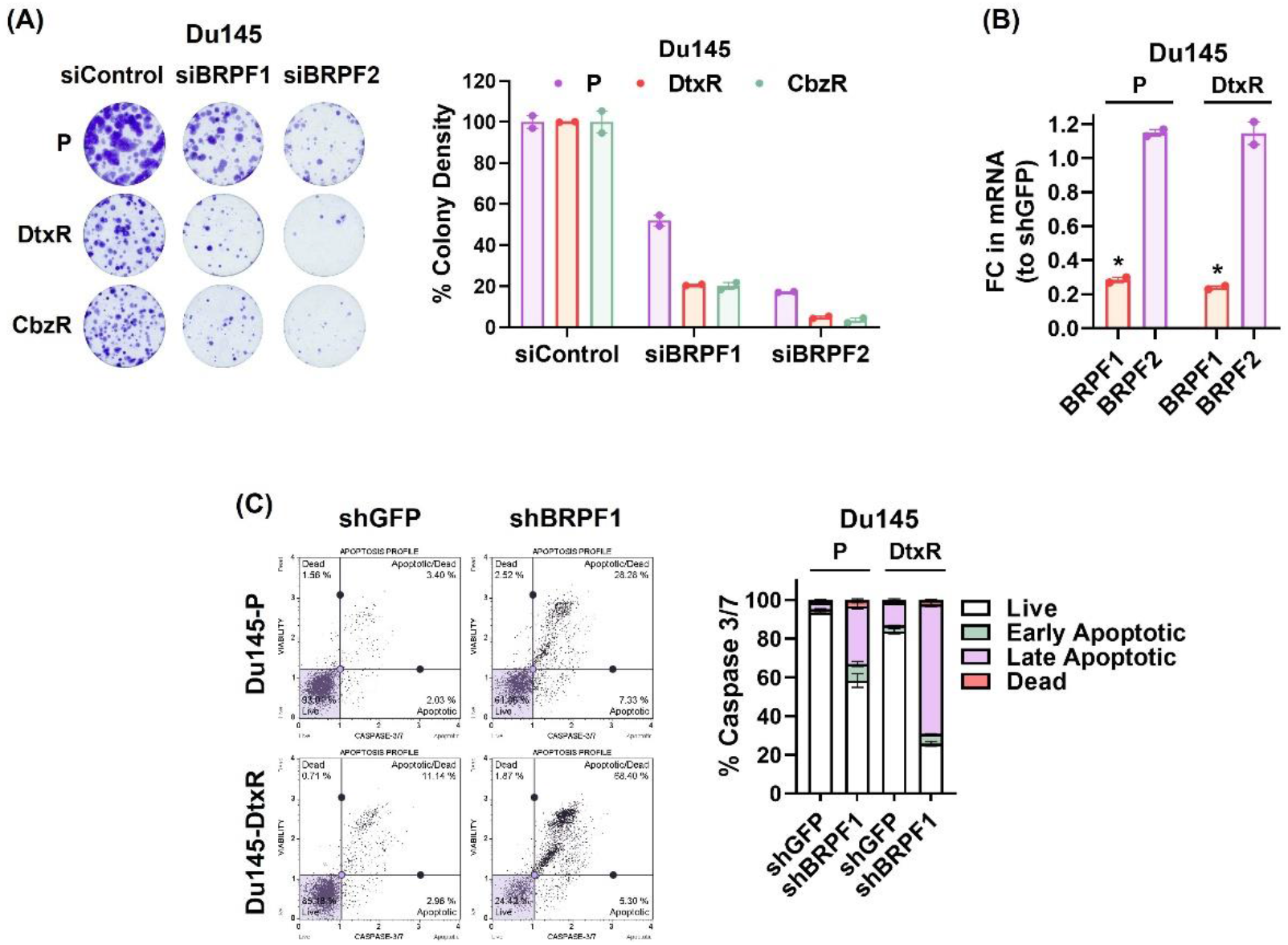
**(A)** Colony forming capability of siRNA treated Du145-P/R cells. Representative images (left panel) and quantifications (right panel) are shown. **(B)** Efficiency and specificity of stable knockdown of BRPF1 in Du145-P and -DtxR cells. After transduction with control (shGPF) and shBRPF1, the expressions of BRPF1 and BRPF2 were detected by qRt-PCR. (*) p < 0.05 indicates significant differences between shGFP and shBRPF1 cells. **(C)** Five days after transduction, cells were analyzed for cell death (Caspase 3/7 activity). Representative plots (left panel) and quantifications are shown (right panel).

**Sup. Figure 20.**
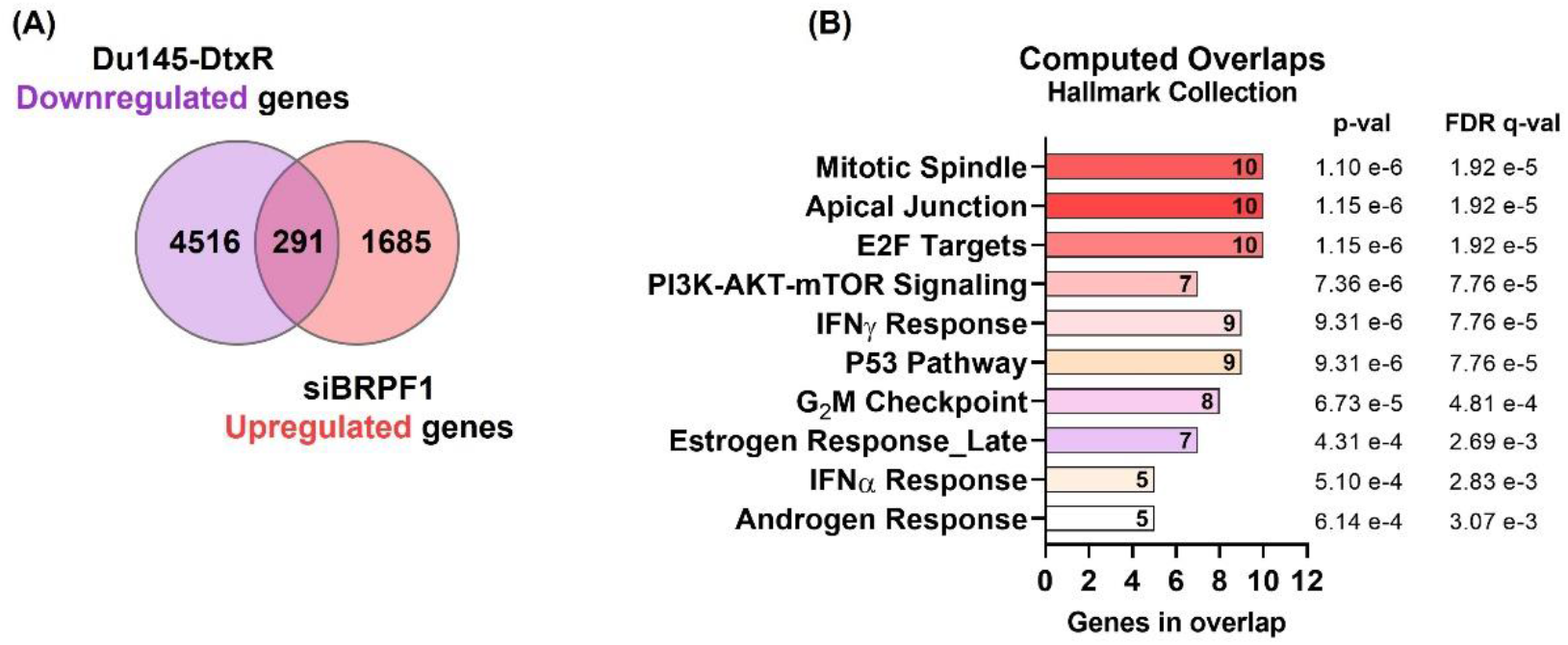
**(A)** Venn diagram showing the number of genes (intersection, 291) whose expression increased after silencing of BRPF1 among genes with decreased expression in Du145-DtxR cells (vs Du145-P). **(B)** Computed overlaps of the 291 genes in the Hallmark Collection of GSEA (MSigDB) database.

**Sup. Figure 21.**
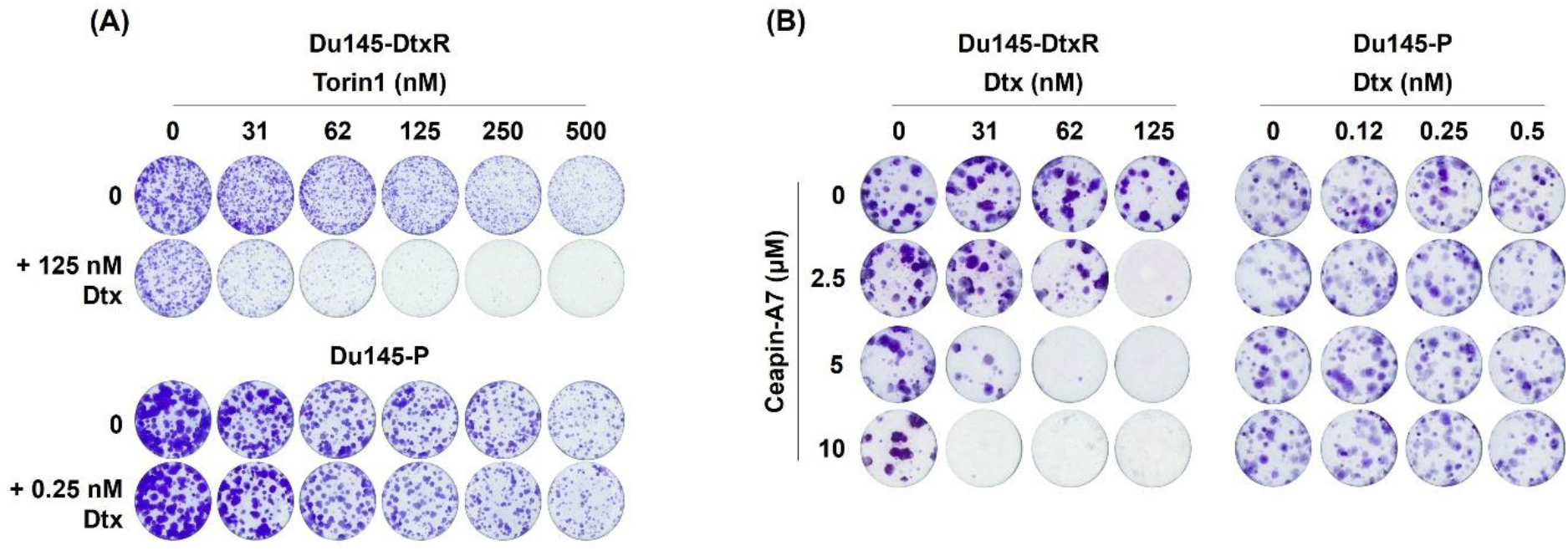
Clonogenic images were obtained by treating Du145-P and -DtxR cells with Torin1 (mTORC1/2 inhibitor, 8-500 nM) and Ceapin-A7 (ATF6α inhibitor, 2.5-10 µM) for 72 h and the colony formation ability was analyzed 10-15 days after drug exposure.

**Sup. Figure 22.**
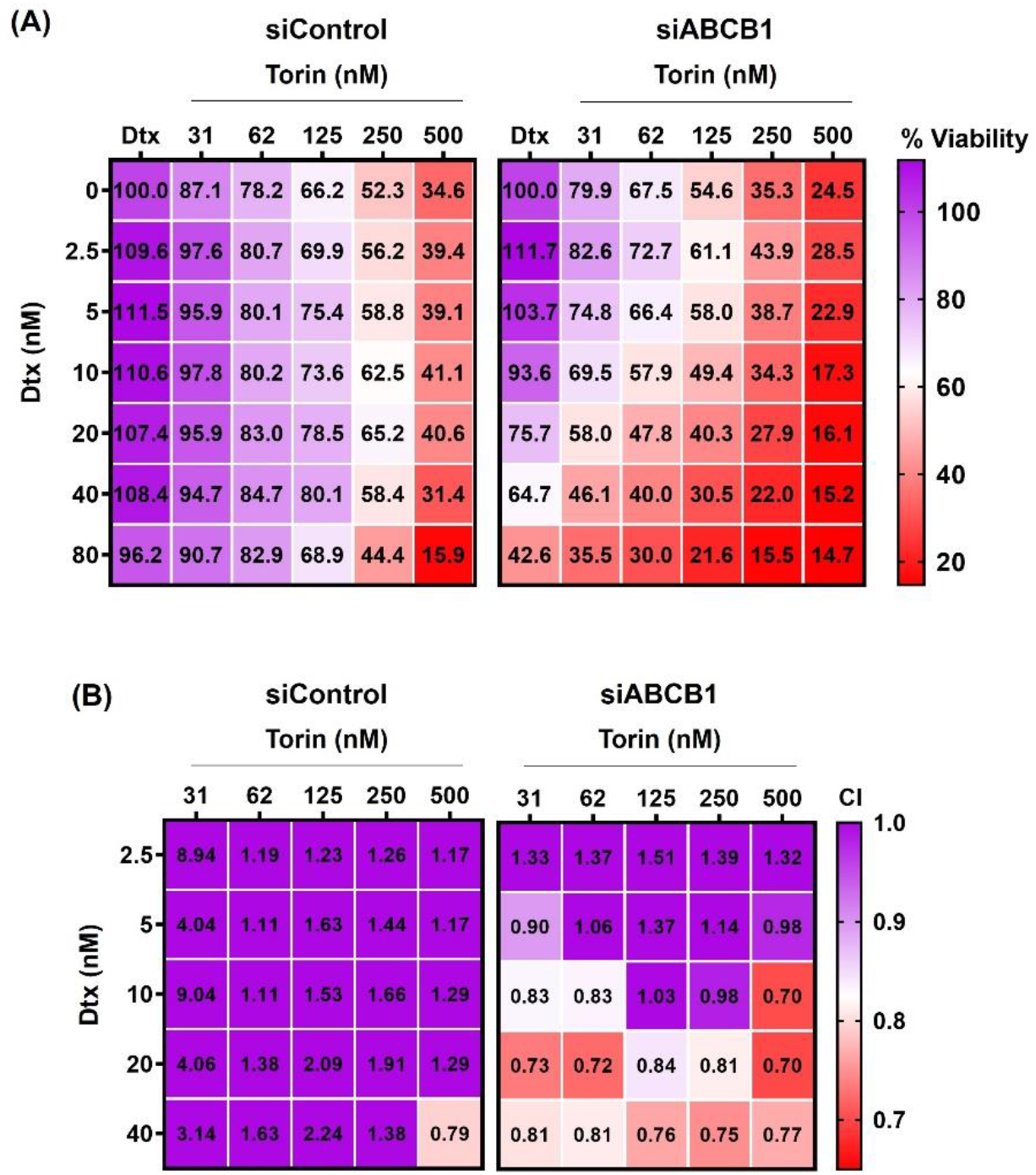
**(A)** The Dtx response of Du145-DtxR cells was assessed after combining ABCB1 knockdown with Torin1 treatment (an mTORC1/2 inhibitor, 31-500 nM). The CTG viability assay was utilized, and the results were represented as a heat map. **(B)** Heat map representation of the Combination Index (CI) values, with red color indicating a synergistic effect. CI was calculated using the CalcuSyn software.

**Sup. Figure 23.**
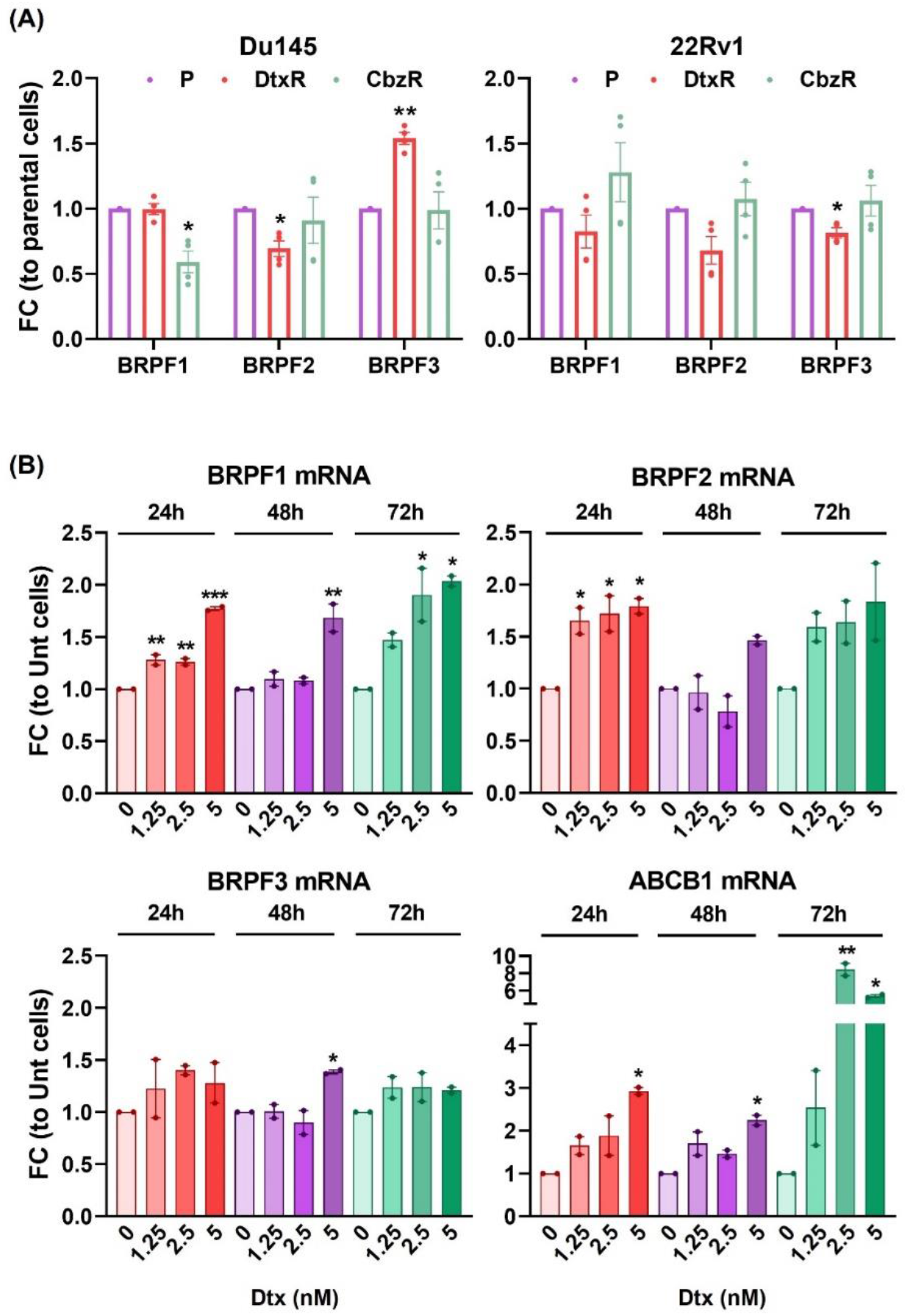
**(A)** The expression profiles of BRPF genes were analyzed by qRt-PCR, comparing taxane resistant cells to their parental counterparts. The data expressed as mean ± SEM from two biological replicates, each performed in duplicate. **(B)** The expression of BRPF genes in Du145-P cells was determined by qRt-PCR following Dtx treatment (1.25-5 nM) at the indicated time points. The data expressed as mean ± SEM. Statistical significance denoted as (*) p < 0.05, (**) p < 0.01, and (***) p < 0.001.

